# The emergence of goblet inflammatory or ITGB6^hi^ nasal progenitor cells determines age-associated SARS-CoV-2 pathogenesis

**DOI:** 10.1101/2023.01.16.524211

**Authors:** Maximillian Woodall, Ana-Maria Cujba, Kaylee B. Worlock, Katie-Marie Case, Tereza Masonou, Masahiro Yoshida, Krzysztof Polanski, Ni Huang, Rik G. H. Lindeboom, Lira Mamanova, Liam Bolt, Laura Richardson, Samuel Ellis, Machaela Palor, Thomas Burgoyne, Andreia Pinto, Dale Moulding, Timothy D. McHugh, Aarash Saleh, Eliz Kilich, Puja Mehta, Chris O’Callaghan, Jie Zhou, Wendy Barclay, Paolo De Coppi, Colin R. Butler, Heloise Vinette, Sunando Roy, Judith Breuer, Rachel C. Chambers, Wendy E. Heywood, Kevin Mills, Robert E. Hynds, Sarah A. Teichmann, Kerstin B. Meyer, Marko Z. Nikolić, Claire M. Smith

## Abstract

Children infected with SARS-CoV-2 rarely progress to respiratory failure, but the risk of mortality in infected people over 85 years of age remains high, despite vaccination and improving treatment options. Here, we take a comprehensive, multidisciplinary approach to investigate differences in the cellular landscape and function of paediatric (<11y), adult (30- 50y) and elderly (>70y) nasal epithelial cells experimentally infected with SARS-CoV-2. Our data reveal that nasal epithelial cell subtypes show different tropism to SARS-CoV-2, correlating with age, ACE2 and TMPRSS2 expression. Ciliated cells are a viral replication centre across all age groups, but a distinct goblet inflammatory subtype emerges in infected paediatric cultures, identifiable by high expression of interferon stimulated genes and truncated viral genomes. In contrast, infected elderly cultures show a proportional increase in ITGB6^hi^ progenitors, which facilitate viral spread and are associated with dysfunctional epithelial repair pathways.

**Graphical Abstract:** 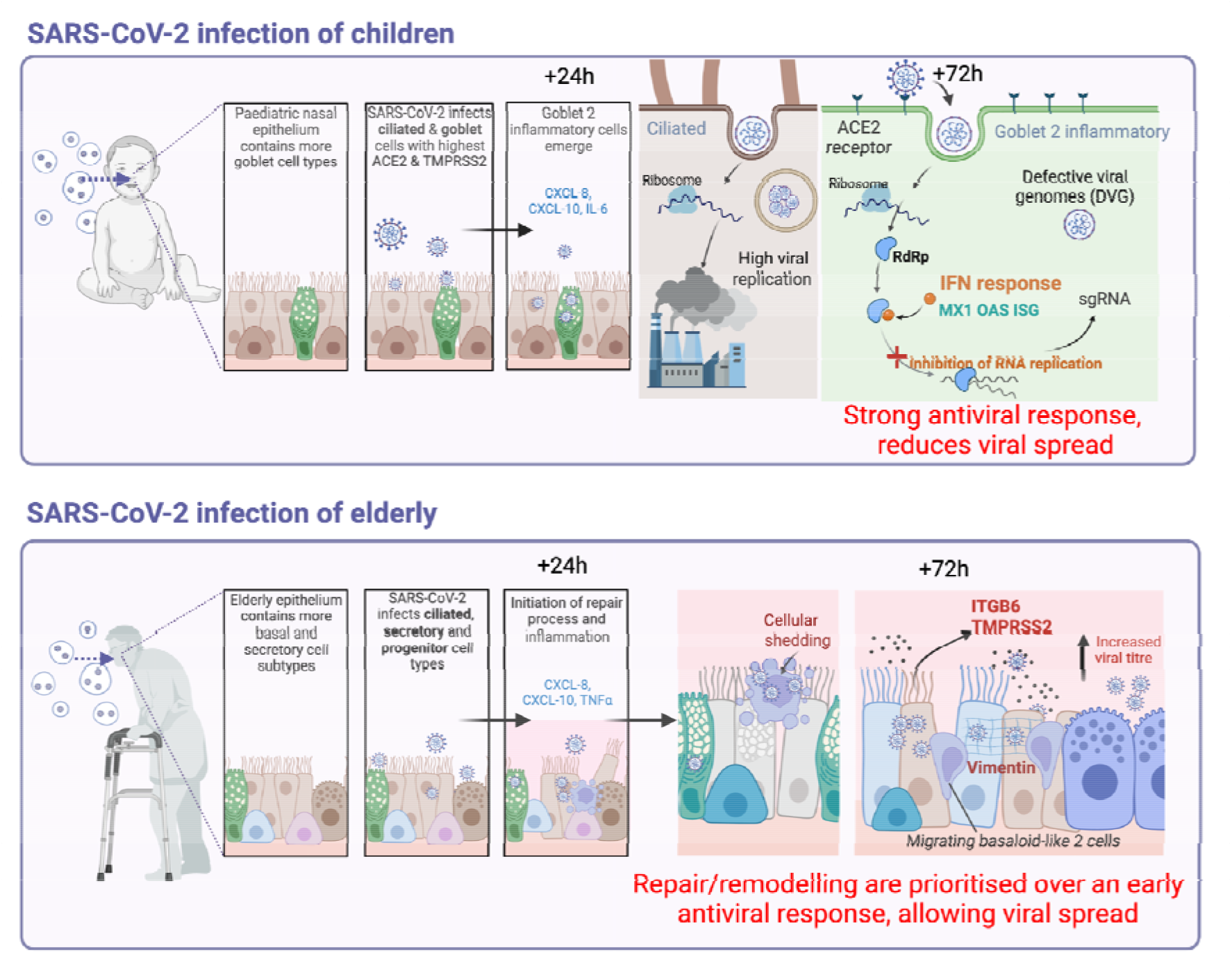

## Introduction

Chronological age remains the single greatest risk factor for COVID-19 mortality, despite the availability of effective vaccines. Children infected with severe acute respiratory syndrome coronavirus 2 (SARS-CoV-2) rarely develop moderate or severe disease^1–4^ whilst the risk of mortality in infected people over 85 years old is currently as high as 1 in 10^5–7^. The nasal epithelium is the first line of defence against inhaled pathogens and nasal epithelial cells (NECs) are the primary target of SARS-CoV-2 infection^8–11^. Infection of upper airway cells^12–14^ can progress distally, leading to diffuse alveolar injury and long-term complications such as the development of lung fibrosis^15, 16^.

It was initially thought that increased COVID-19 severity in the elderly may be due to greater availability of the angiotensin converting enzyme 2 (ACE2) receptor and the host transmembrane serine protease 2 (TMPRSS2) that aid viral entry to NECs compared to children. However, the differences in the expression of these host factors between children and adults are uncertain^17–20^. Current evidence suggests that children at least in part are protected from infection by a pre-activated antiviral interferon (IFN) state in the upper airway epithelium^19, 21–23^. Whilst this may partially explain why children are protected, it does not fully explain the gradient of disease risk seen among adults with increasing age. Understanding epithelial infection and repair in response to SARS-CoV-2 infection has the potential to provide novel therapeutic strategies not only to COVID-19, but also to any future emerging respiratory virus threats.

Here, we investigated the effects of early SARS-CoV-2 infection on human NECs from children (0-11 years old), adults (30-50 years old) and, for the first time, elderly (>70 years old) individuals. NECs were differentiated in air–liquid interface (ALI) cultures. ALI cultures were mock- or SARS-CoV-2-infected for up to 3 days to examine cell-intrinsic differences in function, viral replication, gene and protein expression (**Fig. 1a**). We reveal age-specific responses, with a strong IFN response in infected paediatric goblet inflammatory cells, and the appearance of elderly basaloid-like cells that sustain viral replication and are associated with fibrotic signalling pathways.

**Figure 1.**
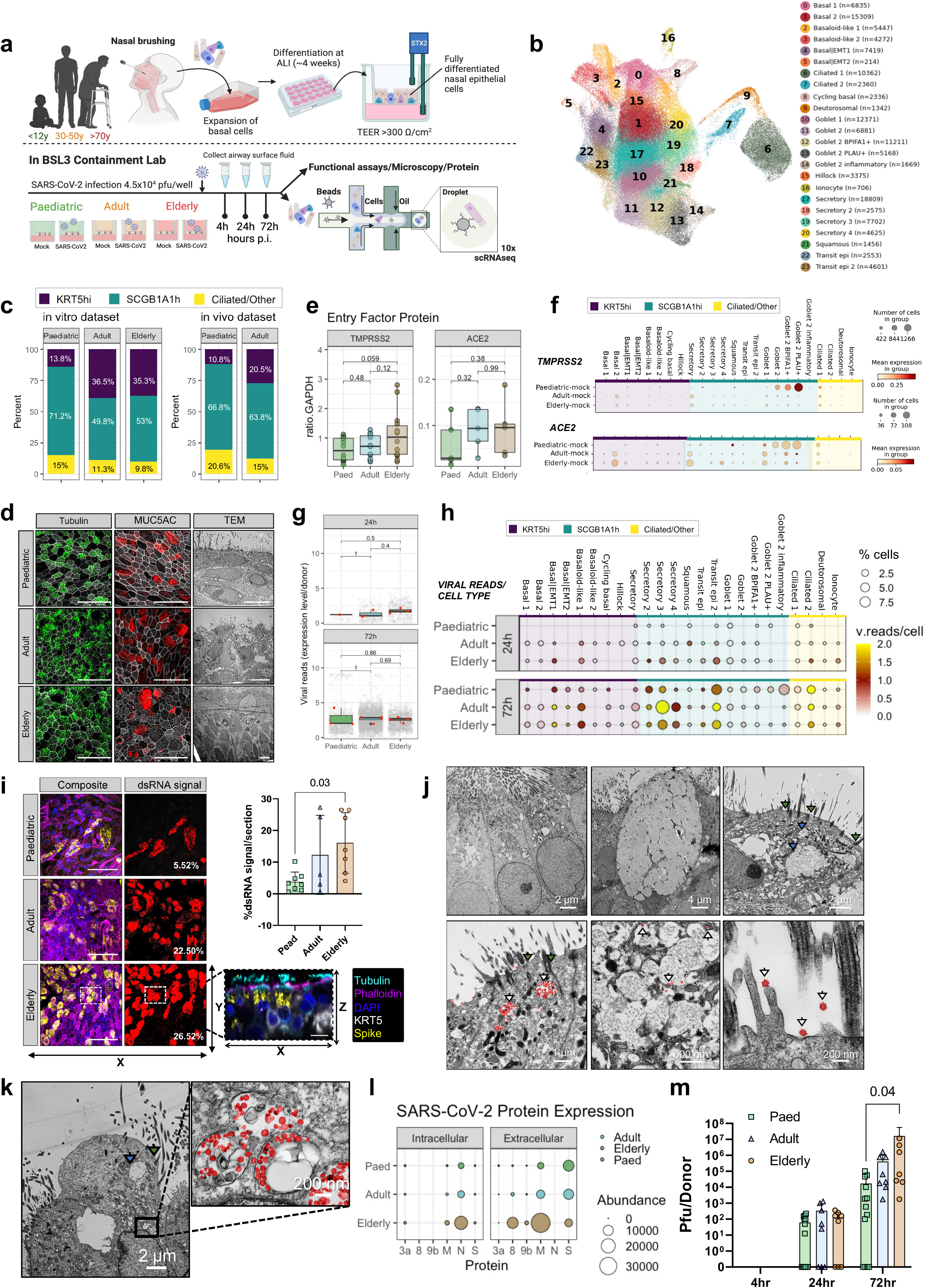
Characterisation of SARS-CoV-2 infected NEC cultures from different age groups. **(a)** Schematic of method and model used to study SARS-CoV-2 infection of paediatric (P), adult (A) and elderly (E) nasal epithelial cells. Human primary nasal epithelial cells (NECs) were cultured on transparent PET culture membrane inserts. NECs were matured at an air-liquid interface (ALI) for 28 days before infection with SARS-CoV-2 England/2/2020 for up to 72 hours with matched mock infections. Samples were collected at 4, 24 and 72 hours post-infection (p.i) for analysis via functional assays, microscopy, proteomics and single cell RNA sequencing (scRNAseq). **(b)** Uniform manifold approximation and projection (UMAP) visualisation of annotated airway epithelial cells. Cell numbers per cell type are shown in parentheses. **(c)** Percentage of new annotation airway epithelial cells in respect to age in baseline (non-infected) ALI cultures and following label transfer to an *in vivo* dataset of nasal brushings from age-matched donors from Yoshida et al,. 2022 ^19^(data shown in percentage cells in the three principal cell domains found in age dataset). (**d**) Representative maximum intensity z-projections of confocal images (left and central) and transmission electron micrographs (right panels) of NEC cultures differentiated at an air–liquid interface and immunolabeled against cilia (tubulin, green) and mucin (MUC5AC, red), with DAPI (blue) and phalloidin (grey) to indicate the nucleus and actin filaments, respectively. Scale bar 10- μm applies to all images in the row. **(e)** SARS-CoV-2 entry factor protein expression per culture type determined by Western blot. ACE2 and TMPRSS2 protein levels normalised to GAPDH (n= P9, A7, E8). **(f)** SARS-CoV-2 entry factor gene expression by scRNAseq. SARS-CoV-2 entry factor gene expression per cell type calculated based upon absolute cell numbers with the average expression of ACE2 and TMPRSS2 indicated by colour. Dot size corresponds to the number of cells expressing ACE2 and TMPRSS2 in respective age groups in the mock condition. **(g)** SARS-CoV-2 RNA viral reads (per cell = grey dots and per donor = red dots) as determined by viral transcript counts (encoding for the full viral genome) per nucleotide per 500 cells (grey dots) or per nucleotide per 500 cells per donor (red dots) within each age group. Pairwise comparisons between donors’ age groups were performed using two-sided Wilcoxon rank-sum tests; P values are shown. Lines in the boxes represent median values. **(h)** SARS-CoV-2 viral reads detected within the scRNAseq dataset (Infected condition only) at 24- (top) and 72-hours post-infection. Shown by cell type and age groups, with dot size and colour indicative of the percentage of cells with detectable viral reads and average reads per cell respectively. **(i)** Representative maximum intensity z-projections of confocal images (left) of NEC cultures differentiated at ALI and immunolabeled against cilia (tubulin, cyan), dsRNA (yellow), basal cells (KRT5, white) with DAPI (blue) and phalloidin (magenta) to indicate the nucleus and actin filaments, respectively. Scale bar is 50-μm. Representation of dsRNA signal alone for each section is indicated in red adjacent to respective maximal projections, the value of spread is given on each panel. Summarised on the bar graph to the right (n=8P, 5A, 6E), subject to one-way ANOVA with Tukey’s multiple comparisons test. A representative orthogonal section is given (bottom right) to indicate location of dsRNA within infected NECs. Transmission electron micrographs of epithelial cell types infected with SARS-CoV-2, with selected areas of interest shown at a higher magnitude for each; **(j)** Ciliated cells (left), Goblet cell (central), Transit (right) and **(k)** Ciliated 2 cell types. Panels show components of interest within each cell type, denoted by arrows: SARS-CoV-2 (white arrows); Cilia (green arrows): secretory mucin granule (blue arrows) and with viral particles false coloured with red to aid visualisation. **(l)** SARS-CoV-2 protein abundance in apical fluid (extracellular) and cell lysates (intracellular) from SARS-CoV-2 infected NECs for 72h p.i as determined by mass spectrometry. Data shown as mean abundance of protein (dot size) and mean fold change in protein abundance per donor from mock infected NECs (colour scale) (n= 5P, 5A, 5E). **(m)** Infectious viral titres in combined cell lysate and apical fluid of SARS-CoV-2 nasal epithelial cells from paediatric (P), adult (A) and elderly (E) donors as determined by plaque assays (n= 13P, 8A, 8E). Subject to two-way ANOVA with Tukey’s multiple comparisons test.

## Results

### Cellular landscape of paediatric, adult and elderly-derived nasal epithelial cultures

We first investigated differences in the cellular landscape of nasal epithelial cultures with age. In total, we generated a single-cell RNA sequencing (scRNAseq) dataset of 139,598 cells, identifying 24 epithelial cell types or states (**Fig. 1b**) based upon key canonical marker genes, including the newly described transit epithelial cells (Yoshida et al., 2022)^19^

(**Extended Data Fig. 1a,b,c** for quality control). We distinguish three broad cell domains: basal (*KRT5^hi^*), secretory (*SCGB1A1^hi^*, *MUC5AC*+) and ciliated (*CCDC40*+) cells (markers in **Extended Fig. 1d**). The first domain includes cell types such as basal, cycling basal, Hillock, Basal|EMT and basaloid-like cells. Two subpopulations of basal cells, named Basal|EMT 1 and Basal|EMT 2, were identified with markers typical of epithelial- mesenchymal transition (EMT), with highest expression of *VIM* and *FN1*; with the latter distinguished by increased expression of *COL1A1* and *COL3A1*. Two similar basaloid-like subpopulations known to be enriched in fibrotic lungs^24^ were also identified, and distinguished from the Basal|EMT populations by higher *ITGB6* and *TGFB1* expression. The second domain includes secretory, goblet and squamous cells (markers in **Extended Fig. 1d**). Several distinct secretory cell populations were identified all expressing secretory proteins such as mucins (*MUC5AC* and *MUC5B*), secretoglobulins (*SCGB1A1*, *SCGB3A1*) as well as genes associated with mucosal defence, including *CAPN13*, *RARRES1* and *SERPINB3*. And lastly the third domain was made up of ciliated cells, which were further subdivided into two clusters based upon the expression of genes known to be involved in cilium organisation including: *OMG*, *PIFO*, *FOXJ1* for Ciliated 1, and *CFAP54A* and *CCDC40* for Ciliated 2 (markers in **Extended Fig. 1d**). A comparison with two existing *in vivo* nasal COVID-19 datasets^19, 25^ provided further confidence in our annotation and highlighted that the ALI model allowed the differentiation of all key epithelial cell types, including rarer cell types such as Ionocytes and Hillock cells (**Extended Data Fig.1e and f**).

Interestingly, in healthy control cultures (non-infected), the proportions of many epithelial cell populations varied with age. Using our cell domain categories, we found that adult and elderly culture datasets contained a greater abundance of basal/progenitor (*KRT5^hi^*) subtypes (**Fig. 1c**), 36.5% and 35.3% respectively, compared to 13.8% in the paediatric cultures. This age-associated difference in basal cell numbers was also seen in an age-matched healthy subset of our previous *in vivo* nasal epithelial dataset^19^ (**Fig. 1c**), using label transfer of our ALI-model cell annotation (**Extended Fig. 1d,e**). Adult and elderly cultures contained the greatest proportion of Basal 1 and Basal 2 cells that were almost absent in paediatric cultures (**Extended Fig. 1g**). Despite this difference in the number of basal cells, all age-groups displayed similar apical differentiation, including mucus (MUC5AC) production and similar levels of cilia (**Fig. 1d** and **Extended Fig. 2a**), with no significant difference in ciliary beat frequency (CBF) (**Extended Fig. 2b**) and no change in cellular motility (measured as the time taken for cells to cover a scratch) with age (**Extended Fig. 2c**). However, ALI cultures from elderly were thicker (mean±SD, 40±18 µm) (**Extended Fig. 1k**) than paediatric cultures (20±10 µm; *p*=0.02) with a distinct spiral morphology typical of ALI cultures (**Extended Fig. 2e**), which was absent in the paediatric cultures. This did not affect the integrity of the epithelial barrier as the trans-epithelial resistance (TEER) was comparable between age groups ranging from mean of 640-450 Ω.cm (**Extended Fig. 2f**).

The most striking difference in the paediatric cultures was the greater abundance of goblet cells, particularly the Goblet 2 subsets (**Extended Fig. 2g**), which appear to represent a shift in cell state from Secretory cells (higher *KRT5*) identified in adult and elderly cultures, to Goblet cells which are higher in *BPIFA1* (see similarity in markers gene expression in **Extended Fig. 1d**). Importantly, whilst there was no overall difference in the protein levels of SARS-CoV-2 entry factors in the cultures with age (**Fig. 1e**), the goblet cell types in the paediatric cultures showed highest expression of *TMPRSS2* (**Fig. 1f**) which is necessary for SARS-CoV-2 spike protein priming ^26^, and *ACE2*, which contains the SARS-CoV-2 spike protein binding site required for viral entry. The highest expression of these markers in the adult and elderly cultures is found in Secretory and Basal 2 cell types (**Fig. 1f**). This suggests a shift in susceptibility to viral infection from goblet to secretory cell types with age. Alternate viral entry factors *BSG*, *CTSL*, *NRP1*, *NRP2* and *FURIN* showed the same trend as *ACE2* and *TMPRSS2* (**Extended Fig. 2g**).

### Increased infectious virus production in SARS-CoV-2 infected elderly cultures

To determine differences in viral replication between age groups, ALI cultures were infected with an early-lineage SARS-CoV-2 isolate (hCoV-19/England/2/2020; 4x10^4^pfu/well (∼multiplicity of infection (MOI) of 0.01 pfu/cell)) for up to 72 hours (h) post infection (p.i). Our preliminary experiments (**Extended Fig. 3a,b**) indicated that over a 5-day infection period, SARS-CoV-2 replication peaked at 72h p.i; therefore, all subsequent investigations were completed prior to this time point. Infectious SARS-CoV-2 particles were measured by performing plaque assays on combined apical washes and cell lysates at 4-, 24- and 72h p.i. We found no significant differences in the level of viral transcripts between age groups at any time point (**Fig. 1g**). However, we found that at 24h p.i SARS-CoV-2 viral reads were detected across more epithelial cell subtypes in adult and elderly cultures (7/24 and 11/24 cell types, respectively) compared to the paediatric cultures, in which viral reads were detected in only 3/24 cell subtypes (**Fig. 1h and Extended Fig. 3c**). At 72h p.i the number of infected cell types increases in all age groups (**Fig. 1h**). This is supported by immunofluorescent analysis at 72h p.i showing greater viral spread (measured as %dsRNA+ signal coverage) in elderly (mean±SD 16.1%±9.5) compared to paediatric cultures (mean±SD 3.8%±3.1) (**Fig. 1i**). Of the cell types expressing viral reads at 72h p.i, we found that Ciliated 2 and Transit epi 2 cells had the highest proportion of viral reads irrespective of age (**Fig. 1h**). The transit epi cell subtype was characterised by the presence of basal (*KRT5*), goblet (*MUC5AC*) and ciliated (*FOXJ1*) markers (**Extended Fig. 1d**) suggesting the potential for both mucus and cilia production. Strikingly, goblet cell types appeared more infected in paediatric cultures, while adult and elderly cultures showed highest viral reads in secretory cell types (**Fig. 1h** and **Extended Fig. 3c,d,e**). Transmission electron microscopy (TEM) demonstrated the presence of viral particles (red) in cells possessing both mucin-containing secretory granules and cilia (**Fig. 1j, k**).

Importantly, we detected a greater apical localisation of the SARS-CoV-2 spike protein (example image **Extended Fig. 3f**), greater abundance of intracellular and apical secreted SARS-CoV-2 proteins (**Fig. 1l**) and higher levels of infectious particles in elderly compared to paediatric cultures, with a significant (*p*=0.04) >800-fold higher titre in elderly (mean±SD: 1.64x10^7^±3.94x10^7^ pfu/well; n=8) compared to paediatric cultures (mean±SD 1.71x10^4^±3.20x10^4^ pfu/well; n=13) at 72h p.i (**Fig. 1m**). These findings support the conclusion that SARS-CoV-2 infected elderly NECs translate more viral protein and generate more replication-competent viruses compared to paediatric cells.

### Emergence of paediatric goblet 2 inflammatory cells in response to SARS-CoV-2, whilst infected elderly epithelium promotes basal cell proliferation

We next profiled the phenotypic effects of infection on epithelial cells, using a diverse combination of live cell microscopy, immunofluorescence staining, proteomics, and gene expression analysis, and compared these across the age groups.

Overall, we found that SARS-CoV-2 infected adult (*p*<0.05, n=5) and elderly (*p*<0.001, n=7) cultures showed a significant decrease in cell culture thickness 72h p.i compared to mock infected cultures that were not seen in SARS-CoV-2 infected paediatric cultures (**Fig. 2a,b, Extended Fig. 4a**). This decrease in culture thickness correlated with a significant decrease in epithelial integrity (TEER) in infected elderly cultures, but not adult or paediatric cultures, at 72h p.i compared to mock infected cultures (*p*<0.03, n=7; **Fig. 2c**). However, there was no significant decrease in E-cadherin, an essential epithelial tight junction protein (**Extended Fig. 4b**). We also detected an increase in basal cell mobilisation in SARS-CoV-2 infected elderly cultures (**Fig. 2d, Extended Fig. 4c,d**) with more non-basal KRT5+ cells detected (∼2-fold) after 72h p.i with SARS-CoV-2 compared to the mock infected cultures (mean±SD, 114±84 compared to 45±41; *p*=0.02) (**Fig. 2d**).

**Figure 2.**
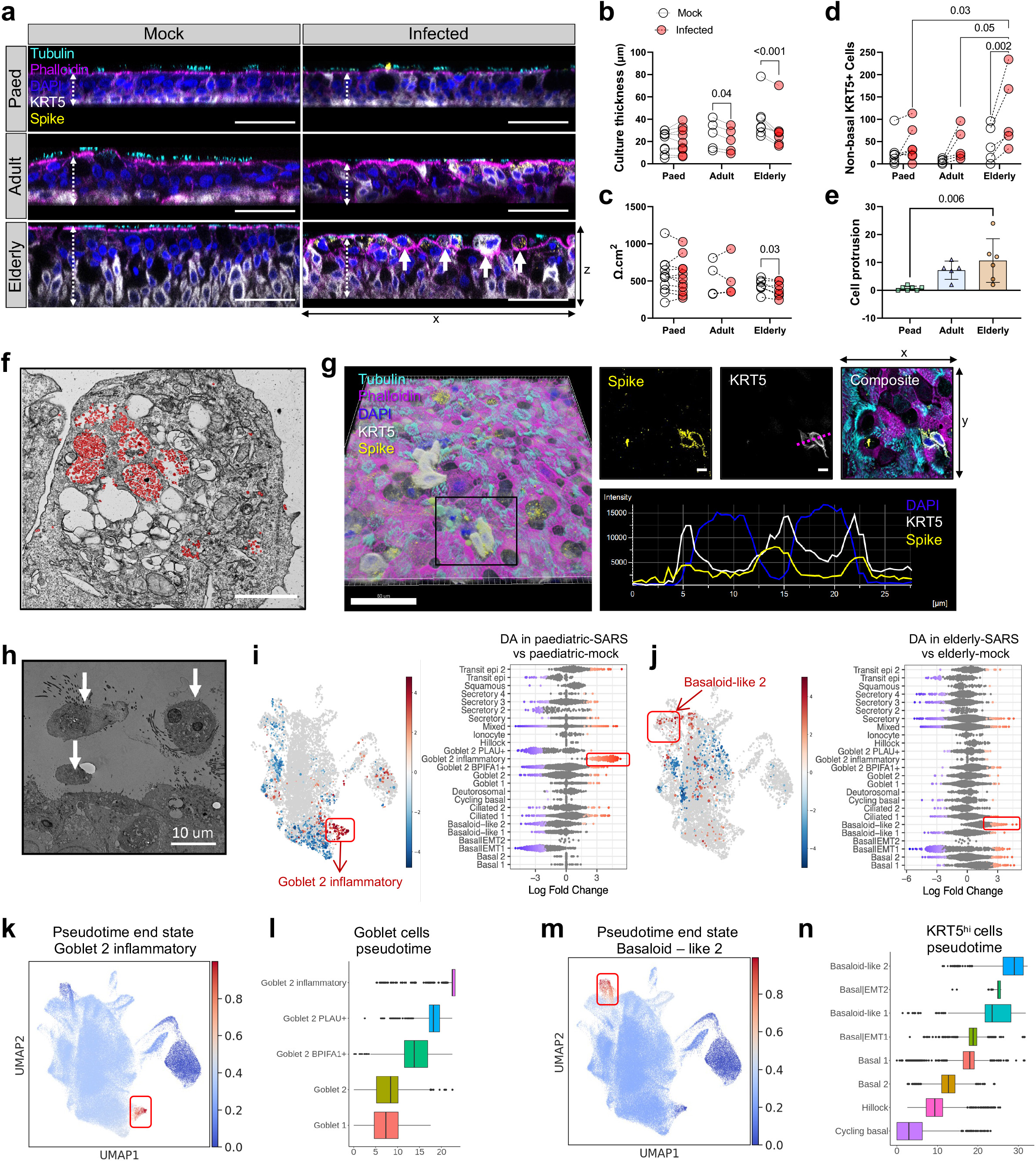
Cytopathology and cellular changes following SARS-CoV-2 infection of NECs. **(a)** Representative orthogonal views of the z-stacks showing the thickness (white dashed arrow) and morphology of fixed paediatric, adult and elderly NECs at 72h p.i mock or SARS- CoV-2 infected. Sections were immunolabeled against cilia (tubulin, cyan), F-actin (phalloidin, magenta) DAPI (blue), SARS-CoV-2 S protein (yellow) and cytokeratin 5 (KRT5+, white). Solid white arrows indicate cells protruding from the apical surface (as quantified in further in (e)) The scale bar represents 50 um. Epithelial thickness **(b)** was further measured and quantified, subject to a two-way ANOVA with Sidak’s multiple comparisons test (n= P9, A5, E7). **(c)** Epithelial integrity, as measured by trans-epithelial electrical resistance (TEER) (Ω .cm2) from 72h p.i. mock or SARS-CoV-2 infected NECs (n= 11P, 4A, 7E). Subject to multiple paired t test. **(d)** Quantification of non-basal KRT+ve cell (e.g KRT+ve cells above and not touching the basal membrane), as a measure of basal cell mobilisation, with age and infection (mock vs infection). Calculated using a cross section of fixed NECs at 72h p.i. (n= P7, A5, E5). Subject to two-way ANOVA with Tukey’s multiple comparison test. See **Extended Fig.5c** for more details for analysis. **(e)** Cell protrusion analysis, calculated by counting the number of nuclei (blue) above apical epithelial membrane (magenta) per section, per donor. For example, as indicated by white solid arrows as in image (a) (n= P7, A5, E6) subject to one-way ANOVA with Tukey’s multiple comparisons test. **(f)** Transmission electron micrograph of protruding epithelial cell type, heavily burdened with virions (red), 72h p.i. with SARS-CoV-2. The scale bar represents 2μm. **(g)** Representative images of immunofluorescence staining for cells that have escaped pseudostratified position and reside above the apical membrane, as stained in (a). Of note here: SARS-CoV-2 spike (yellow) and KRT5 (white). Image 3D-rendered (left) using Imaris (Bitplane) with Blend filter, with a scale bar: 50µm. Size of scale bar for all other images: 5µm rendered in ImageJ (in right bottom panel). A histogram showing distance vs fluorescence intensity for DAPI, KRT5 and SARS-CoV-2 spike staining for a single Z-slice indicated (purple dotted line). **(h)** Transmission electron micrograph of epithelial cell shedding (white arrows) at 72h p.i. with SARS-CoV-2. **(i)** UMAP representation of the results from Milo differential abundance (DA) testing (left plot) and Beeswarm plot (right plot) showing the log fold changes observed when comparing SARS-CoV-2 infected versus mock conditions in the paediatric **(i)** and elderly **(j)**, with a significant enrichment of Goblet 2 inflammatory cells and Basaloid-like 2 cells respectively observed with infection (colour grey is non-significant, colour red is significantly increased, colour blue is significantly decreased at FDR 10%). In the UMAP nodes are neighbourhoods, coloured by their log fold change when comparing SARS-CoV-2 infected versus mock conditions. Node sizes correspond to the number of cells in a neighbourhood and node layout is determined by the position of the neighbourhood index cell in the UMAP. Beeswarm plot shows the distribution of log-fold change across annotated cell clusters when comparing Paediatric-SARS versus Paediatric- mock groups. DA neighbourhoods at FDR 10% are coloured with increased log fold change in red and decreased log fold change in blue. **(k)** Palantir inferred probabilities of cycling basal cells differentiating into Basaloid-like 2 or Goblet 2 inflammatory cells. **(l)** Boxplot showing the distribution of pseudotime within each cluster among Goblet cell subtypes. **(m)** Palantir inferred probabilities of cycling basal cells differentiating into Basaloid-like 2 cells. **(n)** Boxplot showing the distribution of pseudotime within each cluster among KRT5hi cell subtypes.

We also found a significant increase in epithelial escape from pseudostratified culture (cell protrusion) in elderly cultures compared to paediatric at 72h p.i with SARS-CoV-2 (mean±SD, 0.71±0.76 compared to 10.7±7.8; *p*<0.01, n=P7, E6) (**Fig. 2e**); in some instances these cells were heavily burdened with viral particles (**Fig. 2f**) and co-labelled for SARS-CoV-2 spike protein (**Fig**. **2g**). Some protruded cells were shown to completely detach from the pseudostratified epithelium on the apical surface of the culture (**Fig**. **2h, Extended Fig. 4e**).

In terms of ultrastructural changes following infection, we observed mild abnormalities across all age groups, such as endocytosis of cilia basal bodies and sloughing of ciliated cells at 72h p.i using TEM (**Extended Fig. 4f**). However, this did not result in a significant decrease of ciliated cells (a-tubulin protein expression), changes in the ciliary beat frequency or entry factor protein expression over the 72h p.i study period (*p>*0.05) (**Extended Fig. 5a,b,c,d**). These data suggest that, although ciliated cells are a target of SARS-CoV-2 infection, there is no substantial loss of motile cilia within 72h p.i in our model. Proteomic data, including SARS-CoV-2 entry factor expression and mass spectrometry of apical supernatants and cytokine levels are shown in **Extended Fig. 5e**.

Using Milo^27^, a tool to test for differential cell state abundance, we investigated how the cellular landscape of the nasal epithelium changed following infection and whether this differed with age. In paediatric cultures, the airway epithelial cell type composition showed trends of decreasing basal, secretory and goblet cell neighbourhoods (blue) whilst Transit epi 2 and a population of terminally differentiated goblet cells neighbourhoods increased in frequency (**Fig. 2i**). The most striking cellular change following infection was the emergence of Goblet 2 inflammatory cells that were absent from mock-infected paediatric (but not adult or elderly) cultures (**Fig. 2i, and Extended Fig. 5f,g**). The Goblet 2 inflammatory cell type is strongly associated with type I IFN signalling, with higher levels of *CXCL10*, *IFIT1* and *IFIT3* than other goblet cell subtypes (**Extended Fig. 1d**). Whilst goblet inflammatory cells have previously been seen *in vivo*, it is interesting that this inflammatory phenotype is cell intrinsic and independent of immune cells that are not present in our culture system. We later (see next section) explore the impact of this on viral replication and spread.

Infected elderly epithelial cell cultures displayed an increase in basal (*KRT5^hi^*) cell neighbourhoods with SARS-CoV-2 infection compared to mock suggesting an elderly-specific progenitor cell mobilisation (proliferation) following SARS-CoV-2 infection (**Fig. 2j**, adult dataset shown in **Extended Fig. 5h**). We noted a particular expansion of Basaloid-like 2 cell neighbourhoods (**Fig. 2j**). These recently identified cells are poorly described and characterised by markers associated with tissue injury and fibrosis (*ITGB6, ITGB1, ITGAV, ITGB8, VIM, TGFB*) (**Extended Fig. 1d**). In healthy epithelial tissue, including skin and lung, integrin beta 6 mRNA is virtually undetectable^28^, but its expression is considerably upregulated during wound healing^29^, tumorigenesis and the development of fibrosis^30^. The presence of *ITGB6*+ cells are of major interest as they may be involved in exacerbation of disease in the elderly.

To determine the differentiation pathways of these cell types, we used pseudotime trajectory inference and predicted three terminally differentiated end points, including Goblet 2 inflammatory (paediatric) (**Fig. 2k,l**), Basaloid-like 2 (elderly) (**Fig. 2m,n**) and Ciliated 1 cells (see also **Extended Fig. 5i**). Alignment of estimated pseudotimes showed that immediate precursors for the Goblet 2 inflammatory cells are the Goblet 2 PLAU+ cells (**Fig. 2l**), while immediate precursors for the Basaloid-like 2 cells are the Basal|EMT2 cells (**Fig. 2n**).

### Interferon dominates the SARS-CoV-2 response in paediatric cultures leading to truncated viral genomes and fewer infectious viruses

As discussed above, SARS-CoV-2 infection of paediatric epithelial cells led to the emergence of a Goblet 2 inflammatory cell type that was absent in the mock-infected cultures and found in very few numbers in infected older age groups (proportion paediatric = 1455/1578, adult = 90/1578, elderly = 33/1578) (**Extended Fig. 1g**). The Goblet 2 inflammatory cell type has high levels of interferon-stimulated genes (ISGs), particularly *ISG15*, *MX1*, *IF16*, *IFIT3*, *OAS1*, *OAS2*, *ICAM1*, *CXCL10* (**Fig. 3a**) and is strongly associated with type I IFN signalling (**Fig. 3b** and **Extended Fig 6a** and see Gene Set Enrichment Analysis (GSEA) score in **Extended Fig. 6b**), shown to reduce COVID-19 severity^21, 22, 31^. We also applied GSEA to the apical secretome and found biological processes including humoral immune response and activation of immune response (**Extended Fig. 6c,d**). Counterintuitively, in paediatric derived cultures, viral reads were high within the Goblet 2 inflammatory and ciliated cells (**Fig. 1h**); indeed we show colocalisation of MX1 protein with SARS-CoV-2 spike protein (**Fig. 3c**). This suggests that in paediatric cultures the goblet cell types act as a primary cell target for SARS-CoV-2 infection. In support of this, we found the apparent precursor of the Goblet 2 inflammatory subtype, the Goblet 2 PLAU+ cells (**Fig. 2l**) displayed the highest co-expression levels of the serine protease *TMPRSS2* and *ACE2*, that are required for SARS-CoV-2 entry compared to other cell types in the paediatric dataset (**Fig. 3d**). Following SARS-CoV-2 infection, the Goblet 2 inflammatory subtype also displays high levels of these entry factors across the dataset (**Fig. 3d**), indicating that this feature is not lost following differentiation. The gene expression profiles of these subtypes showed that viral replication and the interferon response are distinctive features of the Goblet 2 inflammatory subtype in these cultures (**Fig. 3e**).

**Figure 3.**
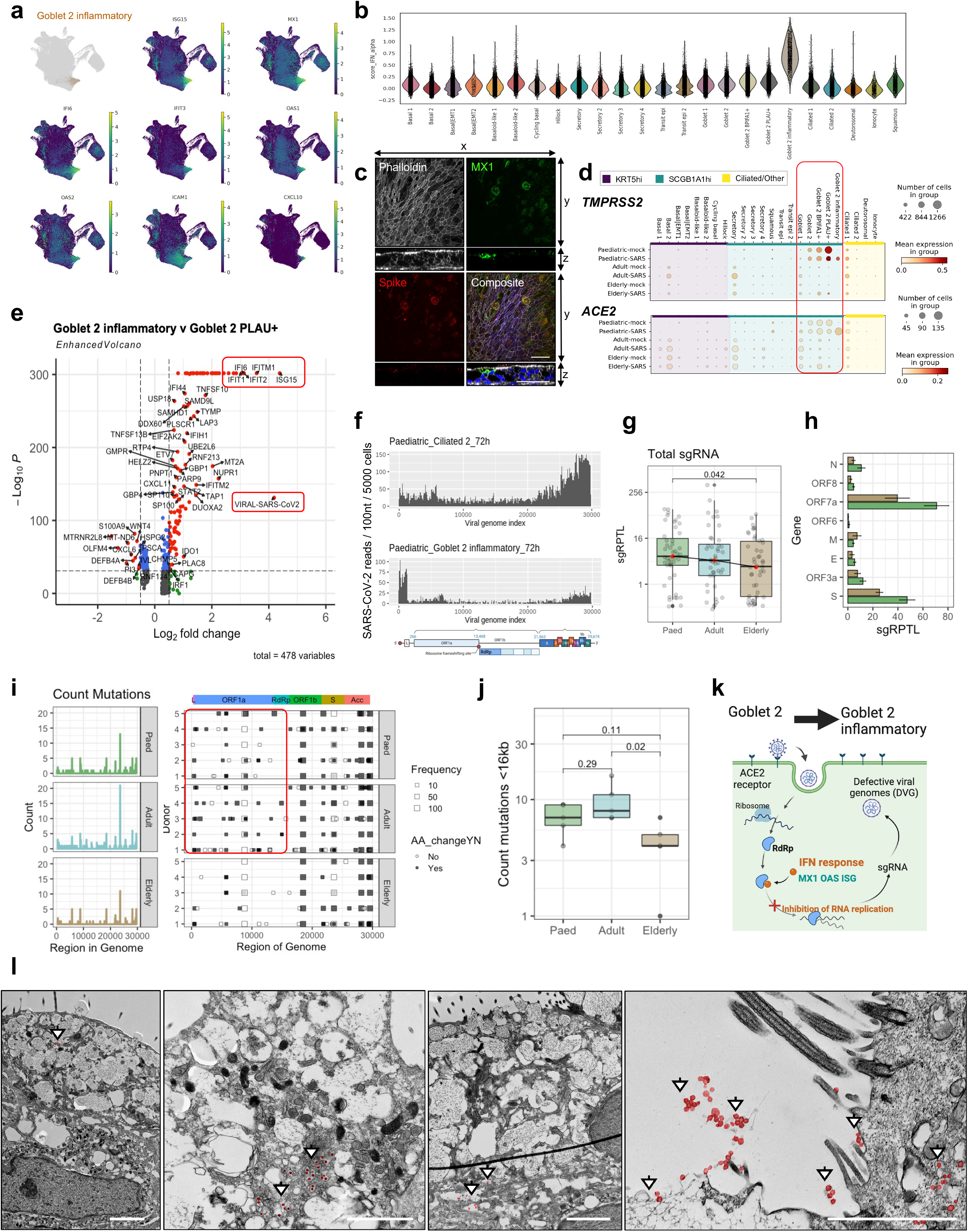
Paediatric Goblet 2 inflammatory cells and truncated viral reads in response to IFN signalling. **(a)** Uniform manifold approximation and projection (UMAP) visualisation of expression of differentially expressed genes in Goblet 2 inflammatory cells. **(b)** Scores of GO term gene signatures for the terms: response to type 1 interferon (GO:0035455 or GO:0034340) across cell types. Scores were calculated with Scanpy as the average expression of the signature genes subtracted with the average expression of randomly selected genes from bins of corresponding expression values. **(c)** Visualisation of MX1 protein expressing cells. Maximum intensity projection images of immunofluorescence staining for F-actin (phalloidin, white), MX1 (green), SARS-CoV-2 S protein (red), with DAPI (blue) in composite image. An orthogonal view of the z-stacks is given in the bottom panel. Example given is a SARS-CoV- 2 infected paediatric culture at 72h p.i. Size of scale bar: 50 μm. **(d)** SARS-CoV-2 entry factor gene expression per cell type calculated based upon absolute cell numbers with the average expression of *ACE2* (top) and *TMPRSS2* (bottom) indicated by colour. Dot size corresponds to Infected number of cells expressing *ACE2* and *TMPRSS2* in respective age groups in the mock (all timepoints) and SARS-CoV-2 (all timepoints) infected condition. **(e)** Volcano plot showing differential gene expression between Goblet 2 inflammatory and their precursor Goblet 2 PLAU+ cells, with a total of 478 variables. Of note were a number of genes associated with a interferon response (e.g. *IFI6*, *IFITM1*, *IFIT1*, *IFIT2* and *ISG15*) and SARS-CoV-2 viral replication (highlighted in red) which were significantly enriched within the paediatric Goblet 2 inflammatory cells. **(f)** Coverage plot of viral reads aligned to SARS-CoV- 2 genome from paediatric ciliated 2 (top) and goblet 2 inflammatory (bottom) cells at 72h p.i. The sequencing depth was computed for each genomic position for each condition. **(g)** Box plot depicting the sgRPTL normalised counts for sgRNA abundances across age groups, with **(h)** showing the distribution of these sgRPTL counts across all genes in paediatric, adult and elderly cultures. Statistical significance is shown or (p ≤ 0.05) is denoted by (*). **(i)** Shows frequency (left) of genomic mutations observed in different regions of the SARS-CoV- 2 genome and (right) the position and if an amino acid change was generated or not from each mutation. Data was generated from 72h p.i with SARS-CoV-2 (n= P5, A5, E5). Bin size is 50 bases. Colour blocks indicate the start coordinates of annotated viral genes. **(j)** Number of genomic mutations occurs <16Kb in genome, shown by age group. **(k)** Hypothesis of SARS-CoV-2 infected Goblet 2 PLAU+ cells becoming protective Goblet 2 inflammatory cell through increased interferon and DVG production. Drawn using Biorender. **(l)** Transmission electron micrographs of Goblet cells 72h p.i. with SARS-CoV-2. The panels show different magnification, the scale bar represents 2 μm. Viral particles are false-coloured in red and indicated with white arrows.

To determine why, despite the high viral reads in paediatric cells, these cultures produce less infectious virions compared to elderly cultures (**Fig. 1m**), we looked at the distribution of RNA reads over the SARS-CoV-2 genome within the Goblet 2 inflammatory and ciliated cell types found in paediatric cultures. Here, we found that viral transcription within the Ciliated 2 cells, which display similar levels of viral reads across the age groups (**Fig. 1h**), showed high viral read expression towards the 3’, underlying that ciliated cells are active sources of viral replication (**Fig. 3f**). However, in the paediatric Goblet 2 inflammatory cells, viral reads were uniquely highest near the 5’ and not towards the 3’, indicating these cells do not replicate virus successfully (**Fig. 3f and Extended Fig. 6e,f,g**). We confirmed that this bias towards the 3’ end is not a technical artefact due to introduction of the spike-in primer to increase detection of viral reads, as SARS-CoV-2 reads were successfully amplified without biassing viral distribution (**Extended Fig. 6h,i,j**). Using deep viral sequencing, we found a greater (*p*=0.042) quantity of non-canonical subgenomic SARS-CoV-2 RNAs (sgRNA) in paediatric and adult samples compared to elderly (**Fig. 3g**), which appeared to be predominantly driven by spike and ORF7a sgRNA (**Fig. 3h**). Whilst canonical sgRNA act as templates for mRNA during SARS-CoV-2 replication, noncanonical sgRNA can result in defective viral genomes (DVGs). These are by-products of the internal recombination processes that form part of normal viral replication. DVGs in other RNA viruses have been associated with antiviral immunity and increased interferon production^32, 33^. In support of this, we also found greater non-canonical (based on the leader sequence at the 5’ end) low frequency and fixed mutations in viral genomes produced by paediatric compared to elderly cultures (**Fig. 3i and Extended Fig. 6k**), particularly before the RNA-dependent RNA polymerase (RdRp) in the viral genome (i.e. <16kB; **Fig. 3j**). Combined with the increased interferon response in these cells, this suggests greater pressure on the virus to mutate in younger cultures compared to elderly cultures. Furthermore, with our earlier data showing a lower infectious viral yield from paediatric cultures (**Fig. 1m**), these data suggest that Goblet 2 inflammatory cells may be directing DVG production, further enhancing the antiviral response in SARS-CoV-2 infected paediatric epithelium (**Fig. 3k**). This is further supported by our ultrastructural (TEM) observation that less viral particles are detected in goblet cells in paediatric cultures compared to neighbouring ciliated cells which were heavily burdened with virus, some of which appear to be within secretory mucin granules (**Fig. 3l and Extended Fig. 7**).

### SARS-CoV-2 infected elderly cultures contain Basaloid-like 2 cells expressing profibrotic and EMT gene signatures

As discussed above, SARS-CoV-2 infection of elderly cultures resulted in a large compositional cell change in the *KRT5^hi^* population, most notably the Basaloid-like 2 cells, which conversely decrease in number with infection in paediatric and adult cultures (**Fig. 4a**). We also show that the Basaloid-like 2 cells are a terminally differentiated *in vitro KRT5^hi^* population (**Fig. 2n**) and that their differentiation is characterised by gene expression of pro-fibrotic factors and markers associated with EMT, including *ITGB6*, *VIM*, *FN1*, *ITGB1*, *ITGAV*, and *TGFB* (**Fig. 4b and Extended Fig. 1d**). ITGAV and ITGB6 and TMPRSS2 were also found to be increased in abundance as proteins in the supernatant of the SARS-CoV-2 infected elderly cultures (**Fig. 4c and Extended Fig. 8a**). ITGB6 and ITGAV are glycoproteins which span the plasma membrane. Therefore their presence in the supernatant/secretome is likely to be from shed cells, exosomes or cellular debris. Vimentin was not found to be upregulated in the secretome, but was found to be significantly upregulated in Elderly cell lysates at 72h p.i with SARS-CoV-2 compared to mock (n=9; *p*<0.05) (**Fig. 4d and Extended Fig. 8b**).

**Figure 4.**
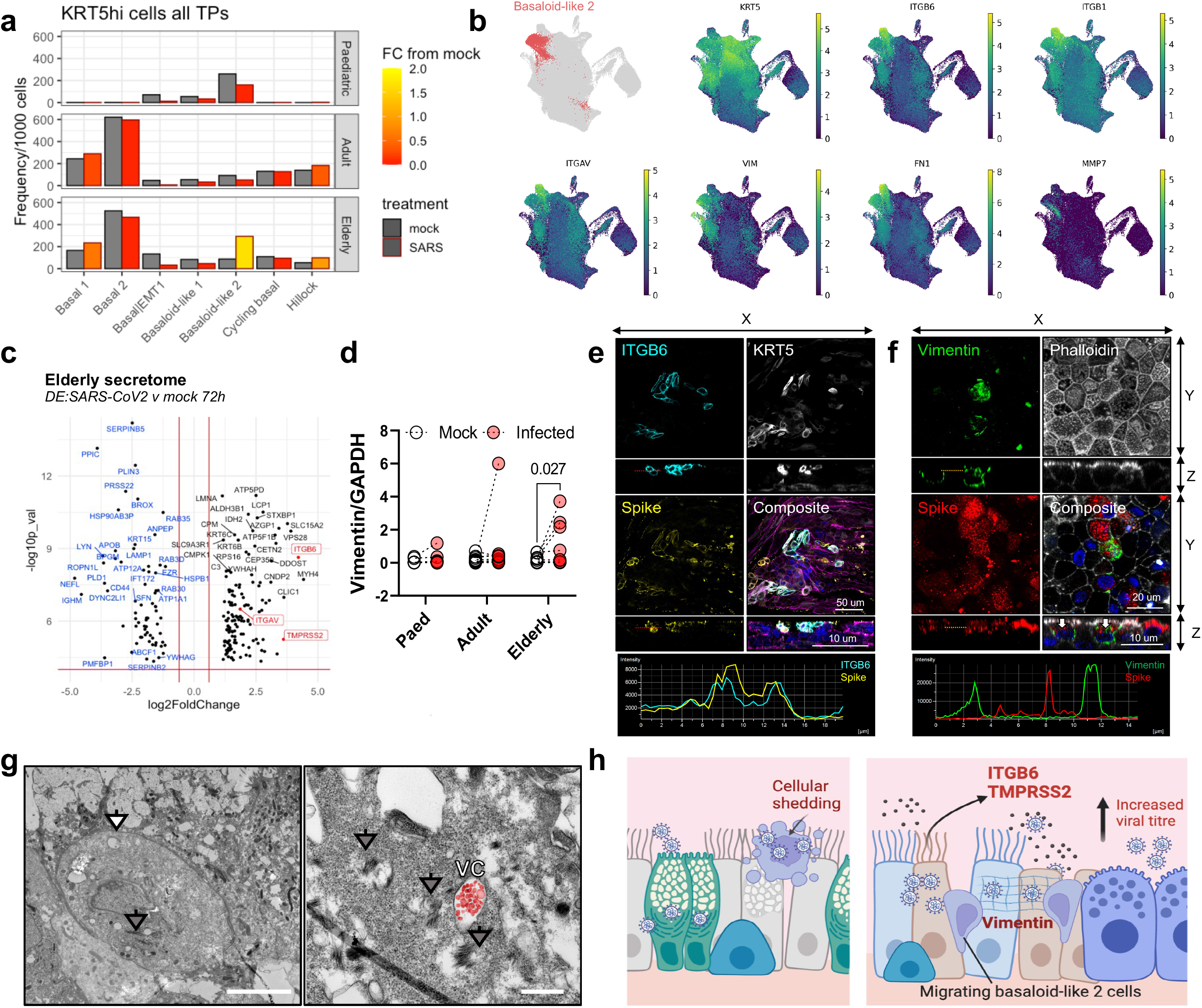
Elderly Basaloid-like 2 cells drive ITGB6 production and enhance viral pathogenesis. **(a)** Frequency of KRT5^hi^ basal airway epithelial cells in mock (black outline) and SARS-CoV- 2 infected (red outline) conditions across all timepoints (4-, 24 and 72- h p.i) in respect to age. Data shown in ratio of cell numbers/per 1000 cells in age dataset, where the colour of the bars indicates fold change (FC) from the matched cell compartment in the mock condition. **(b)** Uniform manifold approximation and projection (UMAP) visualisation of expression of differentially expressed genes in Basaloid-like 2 cells. **(c)** Volcano plot of differential expressed proteins in the apical secretome of mock and SARS-CoV-2 infected cultures that were unique (highly expressed) in the elderly cohort. Blue highlights those that are highly expressed in mock compared to SARS-CoV-2 infection conditions and black enriched with infection, of note; ITGAV, ITGB6 and TMPRSS2 in red. **(d)** Analysis of Vimentin protein levels by western blot normalised to GAPDH (n= P5, A9, E9), subject to multiple paired ratio t test. **(e)** Representative immunofluorescence images of Basaloid-like 2 cell markers in 72h p.i. with SARS-CoV-2. Maximum intensity projection images of immunofluorescence staining in fixed elderly NECs. Left panels: for ITGB6 (cyan), KRT5 (white), SARS-CoV-2 spike (S) protein (yellow) and composite with Phalloidin (magenta) and DAPI (blue). Right panels: Vimentin (green), F-actin (phalloidin: grey), SARS-CoV-2 S protein (red) and composite with DAPI (blue). White arrows annotate the vimentin cage structure around SARS-CoV-2 S protein. Scale differs and scale bars are given on each image**. (g)** Transmission electron micrograph of non-basal KRT5+ve epithelial cell at 72h p.i. with SARS-CoV-2 (white arrow). Cytokeratin bundles are indicated (grey arrows) and viral compartments (VC) containing viral particles false-coloured in red. The scale bars are 5 μm (left) and 0.5 μm (right). **(h)** Hypothesis highlighting an increase in cell protrusion and cell shedding in the elderly, which increase with infection, with cells heavily burdened with viral particles potentially resulting in further spread of infection. This leads to an increase in migrating KRT5+ and ITGB6+ Basaloid-like 2 cells and extracellular matrix (Vimentin cages), which is prioritised over an early antiviral immune response and contributes further to the increase in viral titre seen in the elderly. Drawn using Biorender.

To investigate whether these *ITGB6*+, *VIM*+ and *KRT5*+ expressing cells were permissive to infection and could potentiate viral spread, we analysed cultures by immunofluorescence microscopy, in which we observed ITGB6 protein localisation with SARS-CoV-2 S protein (**Fig. 4e and Extended Fig. 8c,d**) and the formation of a characteristic vimentin cage structure around SARS-CoV-2 S protein (**Fig. 4f and Extended Fig. 8e,f,g**)^34^ in some heavily infected elderly cells at 72h p.i. Using TEM we found rare incidences of non-basal cell types (defined by presence of cytokeratin bundles) burdened with viral compartments (**Fig. 4g and Extended Fig. 8h**) which further suggests that *KRT5*+, *ITGB6*+ and *VIM*+ cells are permissive to infection with SARS-CoV-2.

Interestingly, these Basaloid-like 2 cells show a low level of SARS-CoV-2 transcription (**Fig. 1h**). Therefore, it is likely that infected and damaged Basaloid-like 2 cells may be shed from the epithelium into the secretome (as hypothesised in **Fig. 4h**), supporting our evidence of increased ITGB6 protein (**Fig. 4c**) and increased number of protruding cells found in elderly (**Fig 2e**).

### Dysfunctional repair and ITGB6^hi^ expression enhances viral replication in NECs

To establish the role of the Basaloid-like 2 cells on SARS-CoV-2 pathogenesis we first applied GSEA to the differentially expressed genes in this cell type. The key biological processes associated with this cell type were extracellular matrix and structure reorganisation, response to wounding and a number of migration processes (**Fig. 5a**). Such processes may facilitate viral spread, metastasis and fibrogenic remodelling^35–37^. Furthermore, Basaloid-like 2 cells showed upregulation of alternative SARS-CoV-2 entry receptors *CTSL*, *FURIN*, *NRP1* and *NRP2* (**Extended Fig. 9a** for comparison to other cell types), suggesting they could also be potential targets for infection and spread. This further highlights the upregulation of this cell type as a culprit for poor disease prognosis.

**Figure 5.**
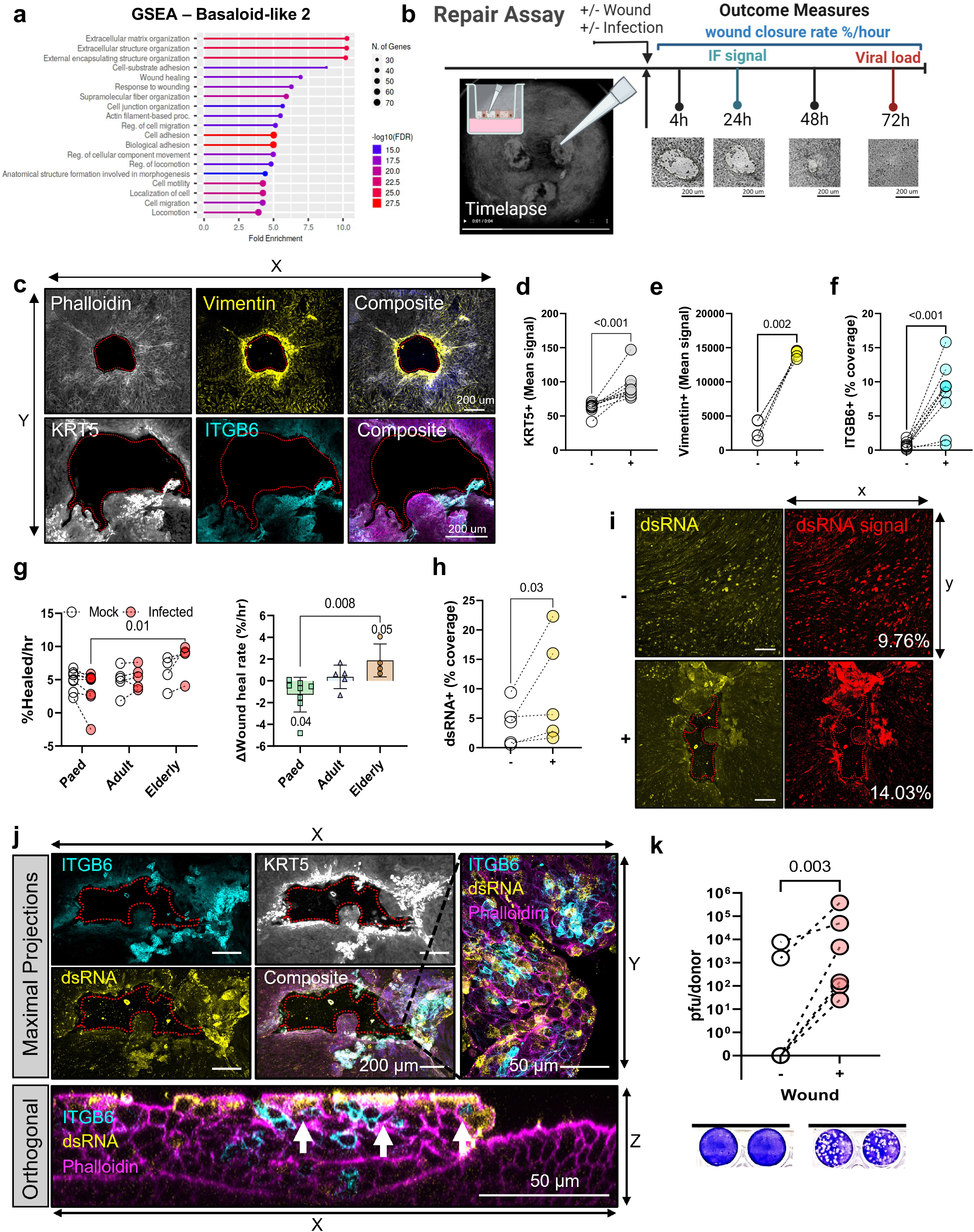
Wound healing response is upregulated in the elderly and associated with increased viral spread. **(a)** Gene Set Enrichment Analysis (GSEA) indicating enriched gene ontology terms for Basaloid-like 2 cells obtained using ShinyGo. **(b)** Schematic to show the different repair assay protocols. **(c)** Representative immunofluorescence images of Basaloid-like 2 cell markers in 24 h post wound NECs. Maximum intensity projection images top panels (left to right): F-actin (Phalloidin; grey), Vimentin (yellow) and composite with DAPI (blue). Bottom panels (left to right): KRT5 (white), ITGB6 (cyan) and composite with F-actin (phalloidin; magenta) and DAPI (blue). Scale bar = 200 µm. Basaloid-like 2 cell markers mean fluorescence signal (MFU) around wound area. Analysis from maximal intensity projections of fixed NECs without (-) and with (+) wounds, 24 h post wounding. KRT5+ve (mean) signal **(d)**, Vimentin +ve (mean) signal **(e)** and ITGB6 +ve % coverage **(f)**. Subject to ratio paired t test (n=6). **(h)** Wound healing rate in NECs from different age groups with mock or SARS-CoV-2 infection. Left panel; Percentage wound closure per hour (%Healed/hr), subject to two-way ANOVA with Sidak’s multiple comparison test. Right panel; The difference in wound closure per hour between mock and SARS-CoV-2 infected from the same donor (n= P8, A5, E4), subject to one-way ANOVA with Tukey’s multiple comparison test. **(i)** dsRNA coverage for NECs irrespective of age group at 72h p.i. Determined by percentage area covered with dsRNA signal (yellow) from maximum intensity projections of fixed NECs. Subject to ratio paired t test (n=5). **(j)** Representative immunofluorescence images of Basaloid-like 2 cell markers ITGB6 (cyan), KRT5 (white), dsRNA (yellow) and F-actin (phalloidin; magenta) in SARS-CoV-2 infected NECs. Maximum intensity projection images from wounded cultures after 24h, shown both as maximal projections (top) and as an orthogonal view (bottom). KRT5 (white) is omitted from composite images, so that overlap of ITGB6 (cyan) and dsRNA (yellow) is apparent (white). Scale differs and scale bars are given on each image. **(k)** Infectious viral titres at 72h p.i in combined cell lysate and apical fluid of SARS-CoV-2 nasal epithelial cells from non-wounded (-) and wounded (+) donors that were previously shown to propagate low levels of infectious particles (<10,000 pfu/donor at 72h p.i.). Infectious viral load in combined apical and cell lysates (pfu/donor) were determined by plaque assays, representative plaque assay wells are shown (bottom). Subject to ration paired t test (n=6).

To investigate whether the Basaloid-like 2 cells gene signature can facilitate viral spread, we stimulated epithelial repair pathways, including mobility, using a wound healing assay (**Fig. 5b**). This assay increased expression of the Basaloid-like 2 cell type markers around the site of the wound, including increased expression of KRT5 protein (**Fig. 5c,d and Extended Fig. 9b,c**) (mean signal±SD, 62.3±8.40 to 94.3±21.1; *p*<0.001, n=9), VIM (**Fig. 5c,e and Extended Fig. 9d,e**) (mean signal±SD, 2887±1378 to 14088±518; *p*<0.002, n=5) and ITGB6 (**Fig. 5c,f and Extended Fig. 9f,g**) (%coverage ±SD, 0.72±0.05 to 8.13±4.73; *p*<0.001, n=9), compared to unwounded cultures (**Fig. 5c**). We further found that the SARS-CoV-2 infected elderly NECs had a significantly (*p*=0.01) faster wound-healing rate (%wound healed/hr, mean±SD, 8.01±2.67% n=4) compared to infected paediatric cultures (3.67±2.78%, n=8) (**Fig. 5g and Extended Fig. 9h**) indicating greater cell mobility which is another indicator of dysfunctional repair processes. SARS-CoV-2 infected elderly cultures also had a significantly faster wound-healing rate compared to their respective mock infected cultures (+1.88%/hr; *p*=0.05, n=4). Interestingly, SARS-CoV-2 infected paediatric NECs, had a significantly slower wound heal rate compared to their respective mock infective cultures (- 1.27%/hr; *p*=0.04, n=8) (**Fig. 5g, right panel and Extended Fig. 9h**).

Importantly, we found that stimulating wound repair correlated with an increase in SARS-CoV-2 infection. The percentage coverage of dsRNA+ve cells (a marker of replicating virus) was increased in wounded cultures (mean±SD, 4.09±3.61% to 9.69±9.04%; *p*=0.03, n=5) (**Fig. 5h and Extended Fig. 9i**), particularly around the site of the wound (**Fig. 5i,j**). We also found that wounding cultures led to an increase in infectious viral particle production from donors that produced low levels of infectious particles (<10^4^ pfu/donor 72h p.i) in the absence of wounding (**Fig 5k**).

## Discussion

In this multi-disciplinary, comprehensive investigation of the early cellular response to SARS-CoV-2 infection of human NECs, we describe several novel age-associated mechanistic differences driving COVID-19 pathogenesis in the elderly. Our key findings are:

● SARS-CoV-2 infected paediatric cultures induce an early, strong type I interferon response emerging from infected Goblet 2 inflammatory cells resulting in truncated viral genomes.
● SARS-CoV-2 infected elderly cultures produce more infectious virus across more epithelial cell subtypes compared to paediatric cultures.
● SARS-CoV-2 infected elderly cultures become thinner, leakier with increased cell shedding compared to controls and paediatric cultures. This leads to an increase in migrating *KRT5*+ and *ITGB6*+ Basaloid-like 2 cells with a gene signature associated with wound repair.
● Wounding cultures prior to infection stimulates expression of the Basaloid-like 2 cell gene signature (*KRT5*+ and *ITGB6*+). This leads to enhanced viral spread and increased infectious viral yield.

### In vitro model and relevance to in vivo datasets

The nose is the primary site of SARS-CoV-2 infection in the human body and fully differentiated primary nasal epithelial cultures are regarded as the gold standard cell culture model to mimic SARS-CoV-2 infection of the airway. However, the response of paediatric and elderly epithelial cultures to SARS-CoV-2 infection has so far been underexplored, particularly at single-cell resolution. Moreover, due to limitations associated with the collection of clinical samples, particularly inaccuracies in the estimated time since infection and sampling after symptom onset, the earliest window of the anti-viral epithelial response remains unresolved.

Recent scRNAseq work found a stereotypical increase in upper airway progenitor basal cell types with age^21, 38, 39^. This finding was supported by our *in vivo* dataset of nasal brushings from healthy adults and paediatric donors^19^, which was further annotated in accordance to our ALI model cell type predictions. In the current study, we also found a greater abundance of basal (*KRT5^hi^*) subtypes in adult compared to paediatric cultures. One of the strengths of our *in vitro* approach is the ability to detect cell-intrinsic differences with age, without confounding factors of intra-individual variations in host immunity. The ability of nasal epithelial cells to recapitulate age-associated differences *in vitro* strongly suggests that these age effects are wired into the epigenome of epithelial cells.

The most striking differences we found across age groups was within the *SCGB1A1^hi^* cell domain, which shifted from the goblet cell state in paediatric cultures to a secretory cell state with age. This is important as these cells expressed the highest amounts of SARS-CoV-2 entry factors (*ACE2* and *TMPRSS2*), and may indicate a shift in viral susceptibility with age. Indeed this was proved to be the case as, excluding ciliated cells, the Goblet 2 and Secretory cells were the primary target of SARS-CoV-2 infection in paediatric and elderly cultures, respectively, containing the highest proportion of these cells expressing viral reads after 72h p.i.

### Cellular innate immunity in paediatric airways

Overall, fewer paediatric cell types expressed viral reads compared to those in the adult and elderly cultures, particularly at 24h p.i. This finding is supported by similar work where adult (29-74y) bronchial epithelial cells (BECs) resulted in rapid infection and progeny production, whilst initial infection of paediatric (5-12y) BECs was limited to few cells^40^. This phenomenon of attenuated infection in children has been attributed to an early IFN response (starting within 24 hours) restricting viral spread^41, 42^. Elderly individuals with severe COVID-19 also displayed type I and II interferon deficiencies, which correlated with SARS-CoV-2 viral load^43^. We suggest that this effect is attributed to the Goblet 2 Inflammatory epithelial cell, the incidence of which deteriorates with age. These cells have the highest viral genome burden and the strongest interferon signature of all epithelial subtypes. This contradiction in high viral burden and high interferon response was explained by our non-standard viral genome index data, indicating truncated viral reads, greater subgenomic RNA and fewer infectious viruses (suggesting more defective viral genomes) compared to SARS-CoV-2 infected elderly cultures. Patient studies have also shown that young children have equivalent or more viral nucleic acid in their upper respiratory tract compared with adults^44^ whilst discrepancies between viral RNA and infectious viral load have also been reported^45–47^. Some animal challenge experiments also support the conclusion that infectious viral production is higher with age^48^.

### Damage and repair in elderly airways

Strikingly, SARS-CoV-2 infection of elderly NECs resulted in epithelial damage and early signs of epithelial repair, including cell migration and proliferation of basal NECs to repopulate damaged areas; events that were not detected in cultures derived from younger age groups. We also detected an increase in ITGAV, ITGB6 and VIM proteins which were attributed to the emergence of Basaloid-like 2 cells. ITGB6 encodes the beta 6 integrin subunit and exclusively partners with the alpha v subunit to form a heterodimeric αvβ6 integrin^49^. This complex is expressed exclusively on epithelial cells but is virtually absent or expressed at very low levels in normal healthy adult epithelium^28, 50^. It is highly upregulated in response to epithelial injury^51^ and is associated with disease progression in the setting of fibrosis and epithelial cancers and is therefore being actively pursued as a major drug target.

Integrins also play a direct role in extracellular signalling and modulating the expression of a number of cytokines and chemokines^52, 53^, including and most notably, binding to and activating the potent cytokine TGF-β1, which has been widely implicated in the development of fibrosis. This axis has also been implicated in triggering EMT^51^, a process with distinct pathological roles in wound healing, tissue regeneration and organ fibrosis and cancer^54^. We therefore hypothesised that SARS-CoV-2 infected NECs undergo reprogramming by these mechanisms in an age-dependent manner and these processes contribute to COVID-19 pathogenesis by delaying disease resolution and enhancing viral spread.

### Disease pathogenesis

Dysregulated epithelial repair processes have previously been identified in the pathogenesis of respiratory infections^55^. We hypothesise that the emergence of the Basaloid-like 2 cell type in SARS-CoV-2 infected elderly NECs drives epithelial-mesenchymal transition (EMT) repair pathways. EMT is a key developmental pathway, which allows for terminally differentiated epithelial cells to dedifferentiate and acquire a mesenchymal-like identity. SARS-CoV-2 and influenza infections^56^ have been previously shown to induce EMT in cell lines and airway epithelia^57,58,59^. However, its role within viral infection, particularly in regards to age, is poorly understood. We show that SARS-CoV-2 infection of elderly cultures results in flattening of epithelial tissue, a decrease in transepithelial resistance, and increase in cell shedding, which are functional markers of the EMT process^60^ and crucial steps in disease progression^61–66^.

### Viral spread

One explanation for increased viral spread in elderly samples could be the direct interaction of the virus with ITGB6, which forms part of the caveolae, a special subcellular structure on the plasma membrane critical to the internalisation of various viruses and respiratory pathogens^51, 67–70^. Recently, a number of *in vitro* studies have suggested that SARS-CoV-2 spike protein interacts directly with integrins^71, 72^, and suggest that they may serve as a viral entry route into non-ACE2 expressing cells, further potentiating infection in the elderly.

In summary, we have shown that SARS-CoV-2 exhibits differential tropism for nasal epithelial cells with age, with preferential infection of paediatric goblet or elderly secretory cell types. Infected paediatric goblet cells mount a robust innate antiviral response to SARS-CoV-2 dominated by interferon, which correlates with truncated viral reads, greater subgenomic viral RNA and less infectious progeny compared to older adult cultures. SARS-CoV-2 infected elderly secretory cells are shed and cultures suffer greater epithelial damage with age. Dysfunctional repair pathways are stimulated and there is an increase in basaloid-like cells that are associated with fibrosis markers and greater viral spread, both features that characterise infection in the elderly. Together these data reveal new insights into age-associated COVID-19 pathogenesis and, for the first time, we reveal how dysfunctional repair enhances SARS-CoV-2 infection in the elderly.

## Materials and Methods

### Participants and ethics

Participants were recruited from five large hospital sites in London, UK, namely Great Ormond Street Hospital NHS Foundation Trust, University College London Hospitals NHS Foundation Trust, Royal Free London NHS Foundation Trust (Royal Free Hospital and Barnet Hospital) and Whittington Health NHS Trust from March 2020 to February 2021. Ethical approval was given through the Living Airway Biobank, administered through the UCL Great Ormond Street Institute of Child Health (REC reference: 19/NW/0171, IRAS project ID: 261511, Northwest Liverpool East Research Ethics Committee). Exclusion criteria for the cohort included current smokers, active haematological malignancies or cancer, known immunodeficiencies, sepsis from any cause and blood transfusions within 4 weeks, known bronchial asthma, diabetes, hay fever, and other known chronic respiratory diseases such as cystic fibrosis, interstitial lung disease and chronic obstructive pulmonary disease. Nasal brushings were obtained by trained clinicians from healthy paediatric (0–11 years), adult (30-50 years) and elderly (≥70 years) donors that tested negative for SARS-CoV-2 (within 24-48 hours of sampling) and reported no respiratory symptoms in the preceding 7 weeks. Brushings were taken from the inferior nasal concha zone using cytological brushes (Scientific Laboratory Supplies, CYT1050). All methods were performed in accordance with the relevant guidelines and regulations.

### Differentiated human nasal epithelial cell culture

Human nasal brushings were collected fresh for this study and immediately placed in a 15-ml sterile Falcon tube containing 4 ml of transport medium (αMEM supplemented with 1× penicillin–streptomycin (Gibco, 15070), 10 ng/ml gentamicin (Gibco, 15710) and 250 ng/ml amphotericin B (Thermo Fisher Scientific; 10746254) on ice. Four matched paediatric nasal brush samples were sent directly for scRNAseq^19^. To minimise sample variation, all samples were processed within 24h of collection and cultured to P1 as previously described^73^. Briefly, biopsies were co-cultured with 3T3-J2 fibroblasts and Rho-associated protein kinase inhibitor (Y-27632) in epithelial cell expansion medium consisting of a 3:1 ratio DMEM:F12 (Gibco, UK; 21765), 1X penicillin/streptomycin and 5% FBS (Gibco; 10270) supplemented with 5 μM Y-27632 (Cambridge Bioscience, UK; Y1000), 25 ng/ml hydrocortisone (Sigma, UK; H0888), 0.125 ng/ml EGF (Sino Biological, UK; 10605), 5 μg/ml insulin (Sigma, UK; I6634), 0.1nM cholera toxin (Sigma, UK; C8052), 250 ng/ml amphotericin B (Gibco, UK; g/ml gentamicin (Gibco, UK; 15710).

Basal cells were separated from the co-culture flasks by differential sensitivity to trypsin and seeded onto collagen I-coated, semi-permeable membrane supports (Transwell, 0.4 µm pore size; Corning, USA). Cells were submerged for 24-48 hours in an epithelial cell expansion medium, after which the apical medium was removed, and the basolateral medium was exchanged for epithelial cell differentiation medium to generate ‘air-liquid interface’ (ALI) conditions. PneumaCult™ ALI medium (STEMCELL Technologies, UK; 05001) and airway epithelial cell growth medium with supplement kit (AECGM, PromoCell, Germany; C-21160) were used for differentiation media following the manufacturers’ instructions. Basolateral media was exchanged in all cultures three times a week and maintained at 37°C and 5% CO_2_. ALI cultures were differentiated in PneumaCult™ ALI medium for four weeks and switched to AECGM for one week prior to the infection experiment.

### Wound healing assay

Mechanical injury of ALI cultures was performed by aspiration in direct contact to the apical cell layer using a P200 sterile pipette tip, creating a wound with a diameter ranging from 750 to 1500 μm. After wounding, the apical surface of the culture was washed with 200 μL PBS to remove cellular debris. The area of the wound was tracked with the aid of time-lapse microscopy with images taken every 60 min at 4x magnification (Promon, AIS v4.6.0.5.). Wound area was calculated each hour using ImageJ. The initial wound area was expressed as 100% to account for variability of wound size. Wounds were considered to be closed when the calculated area fell below 2%, the effective limit of detection due to image processing. Wound closure was calculated as follows: Woundclosure (%)= 100−((Area/Initial Area)∗100). Wound closure (%), plotted as a function of time (h), was used to calculate the rate of wound closure (%/h).

### Virus propagation

The SARS-CoV-2 isolate (hCoV-19/England/2/2020 obtained from PHE) was used in this study. For virus propagation, the African green monkey kidney cell line Vero E6 (a kind gift from The Francis Crick Institute (London, UK) was used. Vero E6 cells were maintained in DMEM supplemented with 5% FCS and 1X penicillin/streptomycin. Cells were fed three times a week and maintained at 37°C and 5% CO2. Vero E6 cells were infected with a multiplicity of infection (MOI) 0.01 pfu/cell in serum-free DMEM supplemented with 1% NEAA, 0.3% (w/v) BSA, and 1X penicillin/streptomycin. The viruses were harvested after 48 hours, aliquoted and stored at -80°C. The titre of virus was determined by plaque assay (see below).

### Viral infection of ALI cultures

After 28 days ALI, differentiated epithelial cultures were rinsed with sterile PBS++ then infected with viral inoculum suspended in PBS++ (4.5x10^4 pfu/ml ∼0.1 MOI) for one hour on the apical compartment at 37°C and 5% CO_2_. The virus inoculums were then removed, and the ALI cultures rinsed and incubated for up to 72h. This time point was chosen as maximum viral replication was observed at day-2-3 in our pilot studies (**Extended Fig. 3a**).

### Infectious viral load quantification by plaque assay

Vero E6 cells were grown to confluence on 24 well plates and then inoculated with serial dilutions of apical supernatant and cell lysates from infected cultures for 1 hour at 37°C and 5% CO_2_. The inoculum was replaced by an overlay medium supplemented containing 1.2% (w/v) cellulose and incubated for 48 hours at 37°C and 5% CO_2_. Plates were fixed with 4% (w/v) PFA for 30 minutes and overlay was aspirated from individual wells. Crystal violet staining was performed for a minimum of 20 minutes, and then plates were washed with water. The number of visible plaques was counted.

### Viral copy number quantification by quantitative real-time PCR

Viral gene quantification was performed on apical wash supernatants from experiments. Samples were lysed in AVL buffer (Qiagen, UK) and stored at -80°C until further processing. The viral RNA extracts were performed using QIAamp viral RNA kit (Qiagen, UK) following the manufacturer’s instructions. 5 μl of extracted RNA samples were determined in one-step RT-qPCR using AgPath-ID one-step RT-PCR (Applied Biosystems, UK) with the following cycle conditions: 45°C for 10 minutes, 95°C for 15 minutes, (95°C for 15 seconds + 58°C for 30 seconds) in total 45 cycles.

Cellular gene quantification was performed with cultured cells collected at the end of the experiments. Cells were lysed in RLT buffer (Qiagen, UK) and extraction was performed using RNeasy mini kit (Qiagen, UK) following the manufacturer’s instructions. Total RNA was converted into cDNA with qScript cDNA supermix (Quantabio, UK) followed by the manufacturer’s instruction. RT-qPCR was performed using TaqMan Fast Advanced Master mix with the following cycle condition: 50°C for two minutes, 95°C for 10 minutes, 95°C for 30 seconds and 60°C for one minute in a total of 45 cycles. The expression was normalised with GAPDH then presented as 2^-(ΔCт) in arbitrary units.

### SARS-CoV-2 genomic sequencing

#### Viral genome read coverage

To visualise the viral read coverage along the viral genome we used the 10X Genomics cellranger barcoded binary alignment map (BAM) files for every sample. We filtered the BAM files to only retain reads mapping to the viral genome using the bedtools intersect tool [52]. We converted the BAM files into sequence alignment map (SAM) files to filter out cells that were removed in our single cell data pre-processing pipeline. The sequencing depth for each base position was calculated using samtools count. To characterise read distribution along the viral genome we counted transcripts of 10 different ORFs: ORF1ab, Surface glycoprotein (S), ORF3a, Envelope protein (E), Membrane glycoprotein (M), ORF6, ORF7a, ORF8, Nucleocapsid phosphoprotein (N) and ORF10.

### Detection of SARS-CoV-2 subgenomic RNAs

Subgenomic RNA analysis was conducted using Periscope (Parker et al., 2021). Briefly, Periscope distinguished sgRNA reads based on the 5′ leader sequences being directly upstream from each gene’s transcription. The sgRNA counts were then normalised into a measure termed sgRPTL, by dividing the sgRNA reads by the mean depth of the gene of interest and multiplying by 1,000.

### Proteomics

#### Mass spectrometry

Paired mock and SARS-CoV-2 infected airway surface fluids from groups of 10 paediatric, adult and elderly cultures were selected for this assay. For mass spectrometry, samples were inactivated with KeyPro™ UV LED Decontamination System (Phoseon Technology, Portland, Oregon USA) before removal from the BSL3. Proteins were precipitated using ice cold acetone. Protein pellets were resuspended in the digestion buffer as previously described and trypsin (Promega, UK) digested to peptides^74^. Peptides were desalted by SPE and separated by reverse phase chromatography on a NanoAquity LC system coupled to a SYNAPT G2-Si mass spectrometer (Waters, Manchester, UK) in a UDMSE positive ion electrospray ionisation mode. Raw MS data were processed using Progenesis QI analysis software (Nonlinear Dynamics, Newcastle-Gateshead, UK). Peptide identification was performed using the Uniprot Human reference proteome, with one missed cleavage and 1% peptide false discovery rate (FDR). Fixed modifications were set to carbamidomethylation of cysteines and dynamic modifications of oxidation of methionine.

### Western blot

Samples were resolved on 4–15% Mini-PROTEAN® TGX™ Precast Protein Gel (Bio-rad: 4561083) with high molecular mass standards 10-250 kDa. Proteins were transferred to a Trans-Blot Turbo Mini 0.2 µm Nitrocellulose membrane in a Trans-Blot Turbo Transfer System (Bio-rad: 1704150). Membranes were blocked in Odyssey® Blocking Buffer overnight. Membranes were then incubated with primary antibodies (dilutions in Odyssey® Blocking Buffer as described) at room temperature for 1 hour. After 3 x 15-minute wash in PBS + 0.1% tween, secondary antibodies (IRDye® 680/800CW IgG) were applied at room temperature for 1 hour. Blots were visualised on an Odyssey Clx imager and quantified with Image Studio™ Lite software.

### Cytokine assay

Apical supernatants were collected by washing the apical surface with 200 μL of PBS. These were snap frozen at -70°C and inactivated with KeyPro™ UV LED Decontamination System (Phoseon Technology, Portland, Oregon USA) in the CL3 laboratory prior to handling them in a CL2 laboratory. Cytokine and chemokine levels was assessed in 25 μL of supernatants using the multiplex BD CBA bead-based immunoassay kits including: IL6: A7; 558276, IL8 (CXCL8): A9; 558277, TNFα: C4; 560112, IFNγ: E7; 558269, IP10 (CXCL10): B5; 558280, IFNα: B8; 560379 and IL10: B7; 558274. Data were acquired using the BD LSRII flow cytometer and concentrations were obtained from a standard curve (provided with the kit). Analysis was performed using the FCAP software v3.0 (BD Biosciences).

### Microscopy

#### Immunofluorescence confocal microscopy

For immunofluorescence confocal imaging, ALI cultures were fixed for microscopy using 4% (v/v) paraformaldehyde for 30 minutes. Cultures were then permeabilized with 0.2% Triton-X (Sigma, UK) for 15 minutes and blocked with 5% goat serum (Sigma, DayUK) for 1 hour prior to overnight staining with primary antibody at 4°C. Secondary antibody incubations were followed the next day for one hour at room temperature. Cultures were then incubated with AlexaFluor 555 phalloidin for an hour at room temperature to stain F-actin, followed by nuclei counterstaining with DAPI (Sigma, UK) for 15 minutes prior to mounting with Prolong Gold Antifade reagent (Life Tech). Samples were washed with PBS-T after each incubation step. All antibodies used in this study are listed in Supplementary Table 2. Images were captured using a LSM710 Zeiss confocal microscope and images rendered and analysed using Fiji/ImageJ v2.1.0/153c^75^ including mean intensity, cell protrusion, culture thickness, % signal coverage, wound area, and pseudocoloring functions. Intensity profiles were plotted with Nikon NIS-Elements analysis module. Imaris software (Bitplane, Oxford Instruments; version 9.5/9.6) was employed for 3D rendering of Immunofluorescence images.

### Transmission electron microscopy

Cultured NECs that were either SARS-CoV-2 infected or non-infected were fixed with 4% paraformaldehyde 2.5% glutaraldehyde in 0.05 M sodium cacodylate buffer at pH 7.4 and place at 4°C for at least 24h. The samples were incubated in 1% aqueous osmium tetroxide for 1h at room temperature (RT) before subsequently en bloc staining in undiluted UA-Zero min at RT. The samples were dehydrated using increasing concentrations of ethanol (50, 70, 90, 100%), followed by propylene oxide and a mixture of propylene oxide and araldite resin (1:1). The samples were embedded in araldite and by leaving at 60°C for 48 h. Ultrathin sections were acquired using a Reichert Ultracut E ultramicrotome and stained using Reynold’s lead citrate for 100 taken on a JEOL 1400Plus TEM equipped with an Advanced Microscopy Technologies (AMT) XR16 charge-coupled device (CCD) camera and using the software AMT Capture Engine.

### Sample preparation for single cell RNA sequencing

The ALI wells were processed using an adapted cold-active protease single cell dissociation protocol^76^, as described below, based upon that previously used^19^ to allow for a better comparison of matched samples included in both studies. The ALI wells carefully transferred into a new 50 mL Falcon tube and any residual transport media carefully removed so not to disturb the cell layer, before 2 mL of dissociation buffer were added to each well ensuring the cells were covered; 10 mg ml^−1^ protease from *Bacillus licheniformis* (Sigma-Aldrich, P5380) and 0.50mM EDTA in HypoThermosol (Stem Cell Technologies, 07935). The cells were incubated on ice for 1 hr. Every 5 min, cells were gently triturated using a sterile blunt needle, decreasing from a 21 G to a 23 G needle. Following dissociation, protease was inactivated by adding 400μl of inactivation buffer (HBSS containing 2% BSA) and the cell suspension transferred to a new 15 mL falcon tube. The suspension was centrifuged at 400*g* for 5min at 4 °C and the supernatant was discarded. Cells were resuspended in 1 mL DTT wash (10mM DTT in PBS)(Thermofisher, R0861) and gently mixed until any remaining visible mucous appears to break down, or for ∼2-4 mins. Spin 400xg for 5 min at 4°C, remove supernatant. Resuspend in 1mL wash buffer (HBSS containing 1% BSA) and centrifuged again. Filter single-cell suspension through a 40-μm Flowmi Cell Strainer. Finally, the cells were centrifuged and resuspended in 30μl of resuspension buffer (HBSS containing 0.05% BSA). Using Trypan Blue, total cell counts, and viability were assessed. 3,125 cells were then pooled together from the 4 biological replicates with corresponding conditions (e.g. all mock viral treatment at 24 hours) and the cell concentration adjusted for7,000 targeted cell recovery according to the 10x Chromium manual (between 700–1,000 cells per μl). The pools were then processing immediately for 10x 5′ single-cell capture using the Chromium Next GEM Single Cell V(D)J Reagent Kit v1.1 (Rev E Guide) or the chromium Next GEM Single Cell 5′ V2 (Dual index) kit (Rev A guide). Each pool was run twice.

For several samples (**Table 2**) 1 μl viral RT oligo (either at 5μM or 100 μM, PAGE) was spiked into the master mix (at step 1.2.b in the 10x guide; giving a final volume of 75μl) to help with the detection of SARS-CoV-2 viral reads. The samples were then processed according to the manufacturer’s instructions, with the viral cDNA separated from the gene expression libraries (GEX) by size selection during step 3.2. Here the supernatant was collected (159 μl) and transferred to a new PCR tube and incubated with 70 μl of SPRI beads (0.6× selection) at room temperature for 5min. The SPRI beads were then washed according to the guide and the viral cDNA was eluted using 30 μl of EB buffer. No changes to the transcriptome were previously observed upon testing the addition of viral oligo^19^, nor were any significant changes observed with an increasing concentration upon comparison, outside of a small increase in the overall number of SARS-CoV-2 reads detected. The RT oligo sequence was as follows: 5′- AAGCAGTGGTATCAACGCAGAGTACTTACTCGTGTCCTGTCAACG-3′

**Table 1.**
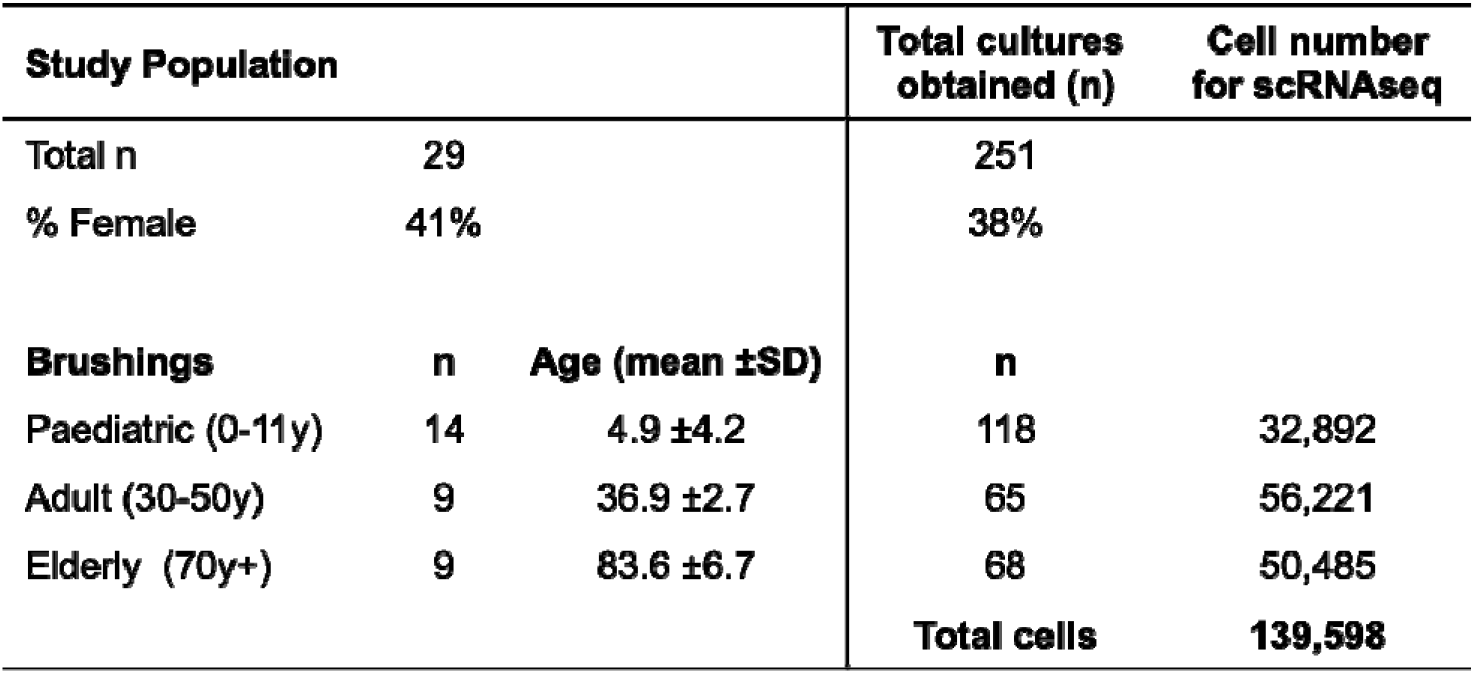
Study population

**Table 2.**

| Lab sample ID | Samples in pool | 10x Kit | Viral Oligo spiked |
| --- | --- | --- | --- |
| ALI1_mock_4h | 276 mock 4h | v1.1 and v2 | NA |
|  | 291 mock 4h |  |  |
|  | 354 mock 4h |  |  |
|  | 357 mock 4h |  |  |
| ALI1_SARS_4h | 276 SARS 4h | v1.1 and v2 | NA |
|  | 291 SARS 4h |  |  |
|  | 354 SARs 4h |  |  |
|  | 357 SARS 4h |  |  |
| ALI1_mock_24h | 276 mock 24h | v1.1 and v2 | NA |
|  | 291 mock 24h |  |  |
|  | 354 mock 24h |  |  |
|  | 357 mock 24h |  |  |
| ALI1_SARS_4h | 276 SARS 24h | v1.1 and v2 | NA |
|  | 291 SARS 24h |  |  |
|  | 354 SARS 24h |  |  |
|  | 357 SARS 24h |  |  |
| ALI1_mock_72h | 276 mock 72h | v1.1 and v2 | NA |
|  | 291 mock 72h |  |  |
|  | 354 mock 72h |  |  |
|  | 357 mock 72h |  |  |
| ALI1_SARS_72h | 276 SARS 72h | v1.1 and v2 | NA |
|  | 291 SARS 72h |  |  |
|  | 354 SARS 72h |  |  |
|  | 357 SARS 72h |  |  |
| ALI2_mock4h | 901 mock 4h | v1.1 | 1uL @ 5µM |
|  | 902 mock 4h |  |  |
|  | 905 mock 4h |  |  |
|  | 907 mock 4h |  |  |
| ALI2_SARS4h | 901 SARS 4h | v1.1 | 1uL @ 5µM |
|  | 902 SARS 4h |  |  |
|  | 905 SARs 4h |  |  |
|  | 907 SARS 4h |  |  |
| ALI3_mock4h | 903 mock 4h | v1.1 | 1uL @ 5µM |
|  | 906 mock 4h |  |  |
|  | 926 mock 4h |  |  |
|  | 927 mock 4h |  |  |
| ALI3_SARS4h | 903 SARS 4h | v1.1 | 1uL @ 5µM |
|  | 906 SARS 4h |  |  |
|  | 926 SARS 4h |  |  |
|  | 927 SARS 4h |  |  |
| ALI2_mock24h | 901 mock 24h | v1.1 | 1uL @ 5µM |
|  | 902 mock 24h |  |  |
|  | 905 mock 24h |  |  |
|  | 907 mock 24h |  |  |
| ALI2_SARS24h | 901 SARS 24h | v1.1 | 1uL @ 5µM |
|  | 902 SARS 24h |  |  |
|  | 905 SARs 24h |  |  |
|  | 907 SARS 24h |  |  |
| ALI3_mock24h | 903 mock 24h | v1.1 | 1uL @ 5µM |
|  | 906 mock 24h |  |  |
|  | 926 mock 24h |  |  |
|  | 927 mock 24h |  |  |
| ALI3_SARS24h | 903 SARS 24h | v1.1 | 1uL @ 5µM |
|  | 906 SARS 24h |  |  |
|  | 926 SARS 24h |  |  |
|  | 927 SARS 24h |  |  |
| ALI2_mock72h | 901 mock 72h | v1.1 | 1uL @ 5µM |
|  | 902 mock 72h |  |  |
|  | 905 mock 72h |  |  |
|  | 907 mock 72h |  |  |
| ALI2_SARS72h | 901 SARS 72h | v1.1 | 1uL @ 5µM |
|  | 902 SARS 72h |  |  |
|  | 905 SARS 72h |  |  |
|  | 907 SARS 72h |  |  |
| ALI3_mock72h | 903 mock 72h | v1.1 | 1uL @ 5µM |
|  | 906 mock 72h |  |  |
|  | 926 mock 72h |  |  |
|  | 927 mock 72h |  |  |
| ALI3_SARS72h | 903 SARS 72h | v1.1 | 1uL @ 5µM |
|  | 906 SARS 72h |  |  |
|  | 926 SARS 72h |  |  |
|  | 927 SARS 72h |  |  |
| ALI3_SARS4h_V2 | 903 SARS 4h | v2 | 1uL @ 100µM |
|  | 906 SARS 4h |  |  |
|  | 926 SARS 4h |  |  |
|  | 927 SARS 4h |  |  |
| ALI3_SARS24h_V2 | 903 SARS 24h | v2 | 1uL @ 100µM |
|  | 906 SARS 24h |  |  |
|  | 926 SARS 24h |  |  |
|  | 927 SARS 24h |  |  |
| ALI3_SARS72h_V2 | 903 SARS 72h | v2 | 1uL @ 100μM |
|  | 906 SARS 72h |  |  |
|  | 926 SARS 72h |  |  |
|  | 927 SARS 72h |  |  |

### Library generation and sequencing

The Chromium Next GEM Single Cell 5′ V(D)J Reagent Kit (V1.1 chemistry) or Chromium Next GEM Single Cell 5 V2 kit (V2.0 chemistry) was used for single-cell RNA-seq library construction for all ALI samples libraries were prepared according to the manufacturer’s protocol (10x Genomics) using individual Chromium i7 Sample Indices. GEX libraries were pooled at a ratio of 1:0.1:0.4 and sequenced on a NovaSeq 6000 S4 Flowcell (paired-end, 150 bp reads) aiming for a minimum of 50,000 paired-end reads per cell for GEX libraries.

### Single-cell RNA-seq data processing

#### Single-cell RNA-seq computational pipelines, processing and analysis

The single-cell data were mapped to a GRCh38 ENSEMBL 93 derived reference, concatenated with 21 viral genomes (featuring SARS-CoV-2), of which the NCBI reference sequence IDs are: <u>NC_007605.1</u> (EBV1), <u>NC_009334.1</u> (EBV2), <u>AF156963</u> (ERVWE1), <u>AY101582</u> (ERVWE1), <u>AY101583</u> (ERVWE1), <u>AY101584</u> (ERVWE1), <u>AY101585</u> (ERVWE1), <u>AF072498</u> (HERV-W), <u>AF127228</u> (HERV-W), <u>AF127229</u> (HERV-W), <u>AF331500</u> (HERV-W), <u>NC_001664.4</u> (HHV-6A), <u>NC_000898.1</u> (HHV-6B), <u>NC_001806.2</u> (herpes simplex virus 1), <u>NC_001798.2</u> (herpes simplex virus 2), <u>NC_001498.1</u> (measles morbillivirus), <u>NC_002200.1</u> (mumps rubulavirus), <u>NC_001545.2</u> (rubella), <u>NC_001348.1</u> (varicella zoster virus), <u>NC_006273.2</u> (cytomegalovirus) and <u>NC_045512.2</u> (SARS-CoV-2). When examining viral load per cell type, we first removed ambient RNA by SoupX^77^. The alignment, quantification and preliminary cell calling of ALI culture samples were performed using the STARsolo functionality of STAR v.2.7.3a, with the cell calling subsequently refined using the Cell Ranger v.3.0.2 version of EmptyDrops^78^. Initial doublets were called on a per-sample basis by computing Scrublet scores ^79^ for each cell, propagating them through an over-clustered manifold by replacing individual scores with per-cluster medians, and identifying statistically significant values from the resulting distribution, replicating previous approaches^80, 81^.

### Quality control, normalisation and clustering

Mixed genotype samples were demultiplexed using Souporcell^82^ and reference genotypes. DNA from samples was extracted as per manufacturer’s protocol (Qiagen, DNeasy Blood and Tissue Kit 69504 and Qiagen Genomic DNA miniPrep Kit) and SNP array derived genotypes generated by Affymetrix UK Biobank AxiomTM Array kit by Cambridge Genomic Services (CGS). Cells that were identified as heterotypic doublets by Souporcell were discarded. Quality control was performed on SoupX-cleaned expression matrixes. Genes with fewer than 3 counts and cells with more than 30% mitochondrial reads were filtered out. Cells with a scrublet score > 0.3 and adjusted *p-*value < 0.8 were predicted as doublets and filtered out. Expression values were then normalized to a sum of 1 × 10^4^ per cell and log-transformed with an added pseudocount of 1. Highly variable genes were selected using the Scanpy function `scanpy.pp.highly_variable_genes()`. Principle component analysis was performed and the top 30 principal components were selected as input for BBKNN^83^ to correct for batch effects between multiplexed pools and kit version and compute a batch-corrected k-nearest neighbour graph. The clustering was performed with the Leiden^84^ algorithm on a k-nearest neighbour graph of a principal component analysis (PCA) space derived from a log[counts per million/100 + 1] representation of highly variable genes, according to the Scanpy protocol^85^. Leiden clustering with a resolution of 1 was used to separate broad cell types (basal, goblet, secretory). For each broad cell type, clustering was then repeated, starting from highly variable gene discovery to achieve a higher resolution and a more accurate separation of refined cell types. Annotation was first performed automatically using Celltypist^86^ model built on the *in vivo* dataset of nasal airways brushes^19^ and secondly using manual inspection of each of the clusters and further doing manual annotation using known airway epithelial marker genes.

### Developmental trajectory inference

Pseudotime inference was performed on the whole object or the basal/goblet compartment using Monocle 3^87, 88^. Briefly, a cycling basal cell was chosen as a “root” cell for the basal compartment, showing the highest combined expression of *KRT5*, *MKI67*, and *NUSAP1* genes. For the goblet compartment, a Goblet 1 cell was chosen as a root, showing the highest combined expression of *TFF3*, *SERPINB3*, *MUC5AC*, *MUC5B*, *AQP5*. Cells were grouped into different clusters using `group_cells()` function, learning the principal graph using `learnGraph()` function and ordering cells along the trajectory using `ordercells()` function. A second pseudotime was inferred with Palantir 1.0.1^89^. The cycling basal “root” cell was determined as above, and an unsupervised pseudotime inference was performed on a Scanpy-derived diffusion map. The five inferred end points were inspected, and three were deemed to be very closely biologically related and replaced with a joint end point with the highest combined expression of *OMG*, *PIFO* and *FOXJ1*. The pseudotime inference was repeated with the two remaining inferred end points and the marker derived one serving as the three terminal states.

### Differential abundance analysis

To determine cell states that are enriched in the SARS-CoV-2 versus mock conditions for the different age groups, we use the Milo framework for differential abundance analysis using cell neighbourhoods^27^. Briefly, we computed k-nearest neighbor (KNN) graph of cells in the whole dataset using the `buildGraph()` function, we assign cells to neighbourhoods using the `makeNhoods()` function and count the number of cells belonging to each sample using the `countCells()` function. Each neighbourhood is assigned the original cluster labels using majority voting. To test for enrichment of cells in the SARS condition versus the mock condition, we model the cell count in neighborhoods as a negative binomial generalised linear model, using a log-linear model to model the effects of age and treatment on cell counts (log-Fold Change, logFC). We control for multiple testing using the weighted Benjamini-Hochberg correction as described in Dann et al.^27^ (SpatialFDR correction). Neighbourhoods were considered enriched in SARS vs mock if the SpatialFDR < 0.1 and logFC > 0.

### Expression signature analysis

To determine the enrichment of basaloid or interferon genes in the annotated clusters, we used the Scanpy function `scanpy.tl.score_genes()` to score the gene signature for each cell. The gene list for computing the basaloid score was composed of the *EPCAM, CDH1, VIM, FN1, COL1A1, CDH2, TNC, VCAN, PCP4, CUX2, SPINK1, PRSS2, CPA6, CTSE, MMP7, MDK, GDF15, PTGS2, SLCO2A1, EPHB2, ITGB8, ITGAV, ITGB6, TGFB1, KCNN4, KCNQ5, KCNS3, CDKN1A, CDKN2A, CDKN2B, CCND1, CCND2, MDM2, HMGA2, PTCHD4, OCIAD2* genes. The gene list for computing the IFN alpha score was composed of the *ADAR, AXL, BST2, EIF2AK2, GAS6, GATA3, IFIT2, IFIT3, IFITM1, IFITM2, IFITM3, IFNAR1, IFNAR2, KLHL20, LAMP3, MX2, PDE12, PYHIN1, RO60, STAR, TPR genes* and the gene list for computing the IFN gamma score was composed of the *OAS3, OASL, OTOP1, PARP14, PARP9, PDE12, PIAS1, PML, PPARG, PRKCD, PTAFR, PTPN2, RAB20, RAB43, RAB7B, RPL13A, RPS6KB1, SHFL, SIRPA, SLC11A1, SLC26A6, SLC30A8, SNCA, SOCS1, SOCS3, SP100, STAR, STAT1, STX4, STX8, STXBP1, STXBP3, STXBP4, SUMO1, SYNCRIP, TDGF1, TLR2, TLR3, TLR4, TP53, TRIM21, TRIM22, TRIM25, TRIM26, TRIM31, TRIM34, TRIM38, TRIM5, TRIM62, TRIM68, TRIM8, TXK, UBD, VAMP3, VCAM1, VIM, VPS26B, WAS, WNT5A, XCL1, XCL2, ZYX* genes, as used in the Yoshida et al, 2022 study.

### Gene set enrichment analysis

Wilcoxon rank-sum test was performed to determine differentially expressed genes (DEGs) between clusters using `scanpy.tl.rank_genes_groups() f`unction. DEGs were further analysed using gene set enrichment analysis via ShinyGO^90^.

### In vivo sub-analysis

Sex- and age-matched healthy adults and paediatric airway samples (n=10 total) were subsetted from our previous dataset^19^ for purposes of label transfer of the *in vitro* cell annotation using CellTypist as described above. Selected sample IDs from the *in-vivo* dataset are shown in **Table 3**. These were selected to match the mean age and range and sex of the current study as the sample collection and processing was conducted in parallel between studies.

**Table 3.** Donor demographics of samples used in the *in vivo* sub-analysis

| Donor ref | Age | Sex | Age Group | Number of cells | Past medical history |
| --- | --- | --- | --- | --- | --- |
| NP-10 | 5y 11month | M | Paediatric | 2882 | Cardiomyopathy |
| NP-14 | 6 weeks | F | Paediatric | 1816 | Supraglottic cyst |
| NP-16 | 3y 11month | F | Paediatric | 5158 | Non-verbal |
| NP-22 | 3 month | F | Paediatric | 405 | Laryngeal cleft |
| NP-31 | 6y 9 month | M | Paediatric | 3046 | unknown |
| AN-9 | 36y | F | Adult | 1359 | non-smoker |
| AN-11 | 38y | F | Adult | 5309 | ex-smoker (3 years) |
| AN-12 | 67y | F | Adult | 348 | non-smoker |
| AN-13 | 55y | M | Adult | 4738 | non-smoker |
| AN-14 | 52y | M | Adult | 837 | non-smoker |

## Acknowledgments

This work was funded by UKRI/ BBSRC (BB/V006738/1) and the NIHR Great Ormond Street Hospital Biomedical Research Centre. We acknowledge funding from Wellcome (WT211276/Z/18/Z and Sanger core grant WT206194). M.Z.N., S.M.J. and K.B.M. have been funded by the Rosetrees Trust (M944, M35-F2) and from Action Medical Research (GN2911). This project has been made possible in part by grants 2017-174169 and 2019- 202654 from the Chan Zuckerberg Foundation. C.M.S is supported by grants from Animal Free Research UK (AFR19-20274), GOSH Children’s charity (COVID_CSmith_017) and the Wellcome Trust (212516/Z/18/Z). M.Z.N. acknowledges funding from a MRC Clinician Scientist Fellowship (MR/W00111X/1) and from the Rutherford Fund Fellowship allocated by the MRC UK Regenerative Medicine Platform 2 (MR/5005579/1). Microscopy was performed at the Light Microscopy Core Facility, UCL GOS Institute of Child Health supported by the NIHR GOSH BRC award 17DD08. The views expressed are those of the author(s) and not necessarily those of the NHS, the NIHR or the Department of Health.

## Author contributions

These authors contributed equally: Maximilian Woodall, Ana-Maria Cujba, and Kaylee B. Worlock.

These authors jointly supervised this work: Sarah A. Teichmann, Kerstin B. Meyer, Marko Z. Nikolić, Claire M. Smith.

M.W., A.M.C., K.B.W., M.Y., K.M.C. designed the study, conducted experiments, analysed data, and reviewed the manuscript. K.B.W. and M.Y. performed 10x and isolated DNA for genotyping. K.P., N.H., and R.G.H.L. assisted with computational analysis, including providing code for viral distribution analysis and customised dotplot functions. A.S. E.K., P.M., C.O., C.R.B., P.d.C., and M.Z.N. recruited patients, and collected nasal brushings and clinical metadata. A.P. and T.B. assisted with transmission electron microscopy image analysis and review of the manuscript. T.M. assisted with flow cytometry experiments and scRNAseq data analysis and review of the manuscript. S.R. and J.B. assisted with viral genomic data analysis and review of the manuscript. W.H., H.V. and K.M. assisted with mass spectrometry data analysis and review of the manuscript. D.M. assisted with microscopy acquisition and reviewed the manuscript. C.M.S., C.O., P.D.C., and C.R.B. provided support through ethics and patient recruitment. S.R. and J.B. oversaw viral genomics analysis and interpretation and review of manuscript. J.Z. and W.B. facilitated collection of preliminary data. T.Mc. provided support in setting up and training for all CL3 work. P.D.C., C.O., W.B, S.A.T, K.B.M, C.R.B, R.E.H, M.Z.N. and C.M.S. conceived the study and oversaw the funding application, contributed to study conception and design and review of the manuscript. S.A.T, R.C., K.B.M, R.E.H, M.Z.N. and C.M.S. oversaw data analysis and interpretation, and the write-up of the manuscript.

## Extended Figure Legends

**Extended Figure 1 Paediatric nasal epithelial cells have less basal cell subtypes compared to adult and elderly nasal epithelial cells but display comparable differentiation markers and SARS-CoV-2 entry factor expression.**

**(a)** Violin plot visualisation of thresholds for quality control steps such as scrublet score and percentage of mitochondrial reads. **(b)** Uniform manifold approximation and projection (UMAP) visualisations showing good integration of donor IDs, donor pool, treatment, age group, 10X Chromium kit version, sex, cell cycle phase and introduced spike-in primer after batch correction (see Methods for more details). **(c)** Boxplot indicating comparison of cell cycle phase states (G1, G2M, S) amongst the three age groups in mock/infected/combined (All) conditions. **(d)** Dot plot visualisation showing marker genes for annotated airway epithelial cell types, with fraction of expressing cells and average expression within each cell type indicated by dot size and colour, respectively. Broad cell domains are colour coded; KRT5^hi^ (purple), SCGB1A1^hi^ (green) and ciliated/other (yellow). Logistic regression based label transfer using Celltypist for the data sets in **(e)** Yoshida et al., 2022 ^19^ and **(f)** Ziegler et al., 2021 ^25^, with fraction of matched cells and average probability score indicated by dot size and colour, respectively **(g)** Numbers of annotated airway epithelial cells in respect to age (data shown in ratio of cell numbers/per 1000 cells in age dataset).

**Extended Figure 2. Physiological differences in paediatric, adult and elderly nasal epithelial ALI cultures.**

Physiological comparison of paediatric (P), adult (A) and elderly (E) ALI cultures as measure by; **(a)** the percentage of +ve -tubulin staining coverage per 225 μm2 section (n=6P, 4A, 5E); **(b)** ciliary beat frequency (CBF)(Hz) (n= 6P, 6A, 7E); **(c)** motility measured by percentage scratch closure over time (n=3P, 3A, 3E) and **(d)** culture thickness (n=9P, 5A, 7E). For (**a-c**) data was subject to one-way ANOVA with Tukey’s multiple comparison test **(e)** Representative light microscope images of whole well scans of ALI cultures depicting the characteristic differences in culture morphology with age (n= 3P, 3A, 3E). **(f)** Comparison of epithelial integrity via trans-epithelial electrical resistance (TEER) (Ω .cm^2^) (n= 12P, 8A, 8E) across the age groups using one-way ANOVA with Tukey’s correction. **(g)** Alternate SARS-CoV-2 entry factor gene expression per cell type calculated based upon absolute cell numbers with the average expression of *BSG, CTSL, NRP1, NRP2* and *FURIN* indicated by colour. Dot size corresponds to the total number of cells expressing alternate viral entry genes in respective age groups in the mock condition.

**Extended Figure 3. Elderly cells replicate more infectious viruses with a greater distribution of viral reads across epithelial subtypes.**

**(a)** Preliminary data showing viral replication over a 5 day period and peaking at 72h p.i. **(b)** Exemplar image of plaque assay used to determine infectious viral load.**(c)** UMAP of airway cells with detected SARS-CoV2 mRNA in each cell type in cultures mock and SARS-CoV-2 infected for all time points and ages combined (≥1 viral UMI per donor following filtering out ambient RNA). **(d)** Pie charts showing the fractions of annotated airway epithelial cells containing viral reads at 4h, 24h and 72h p.i in respect to age. **(e)** Numbers of annotated airway epithelial cells containing viral reads at 24h and 72h post infection (red) in respect to age with total number of cells in each subset (grey) (data shown as cell numbers/per 1000 cells in age dataset). **(f)** Representative orthogonal views of z-stacks from fixed paediatric (top), adult (middle) and elderly (bottom) NECs at 72h p.i with SARS-CoV-2. Sections were immunolabeled against F-actin (phalloidin, grey) DAPI (blue) and SARS-CoV-2 S protein (red). The scale bar represents 10 um.

**Extended Figure 4. Minimal cytopathology following SARS-CoV-2 infection of NECs.**

**(a)** Orthogonal views of the z-stacks showing the thickness and morphology from fixed paediatric, adult and elderly NECs from 3 separate donors at 72h p.i mock or SARS-CoV-2 infected NECs. Sections were immunolabeled against cilia (tubulin, cyan), F-actin (phalloidin, magenta) DAPI (blue), SARS-CoV-2 S protein (yellow) and cytokeratin 5 (KRT5+, white). **(b)** Representative maximal intensity projections (top panels) and orthogonal z sections (bottom panels) of NECs stained for tight junctions (phalloidin) at 72h p.i with mock or SARS-CoV-2, expression of tight junction protein E-Cadherin was determined via western blots and normalised to GAPDH (P5, A5, E5) Subject to one-way ANOVA with Tukey’s multiple comparison test. **(c)** Representative orthogonal views of the z-stacks showing the localisation of KRT5+ve cells (white) from example air–liquid interface cultures (summary in Fig. 2d). KRT5+ve cells above the horizontal yellow line are classified as non-basal -layer (not in contact with basement membrane), F-actin (phalloidin, magenta) stain references apical membrane and tight-junctions. Each section is 225 um in width. **(d)** Maximal projections of the z-stacks showing the localisation of KRT5+ve cells (white) from above and below the horizontal yellow line (classified as non-basal -layer) from example air– liquid interface cultures at 72h p.i with mock or SARS-CoV-2. **(e)** Transmission electron micrograph of epithelial cell shedding at 72h p.i with SARS-CoV-2. **(d)** Transmission electron micrograph of epithelial cell damage at 72h p.i with SARS-CoV-2. Virions (white arrows), viral compartments (VC), Cytopathology including endocytosed cilia basal bodies, loss of tight junctions (grey arrows).

**Extended Figure 5. Cytopathology and changes in cell types**

**(a)** Cilia coverage for each age group at 72hpi Mock and SARS-CoV-2. (left) Representative Immunofluorescent images from 72h p.i NECs with mock and SARS-CoV-2 condition stained for a-tubulin (cyan). Percentage area covered (right) with αTubulin+ve signal (cyan) from maximum intensity projections of fixed NECs using threshold analysis (red) in ImageJ. Summary of cilia coverage (right) (n= P5, A4, E5). Subject to one-way ANOVA with Tukey’s multiple comparison test. For 72h p.i (mock and SARS-CoV-2 conditions) NECs were compared between age groups, looking at **(b)** cilia beat frequency (Hz) (n= P8, A8, E8); **(c)** α-Tubulin protein expression (n= P11, A8, E7) and **(d)** SARS-CoV-2 entry factor protein expression (ACE2, TMPRSS2 and short ACE2). Protein levels were determined via Western blot and normalised to GAPDH (n= P6-8, A5-9, E5-8). **(e)** Volcano plot of the apical secretome showing differential expressed proteins per age group between mock and SARS-CoV-2 infected cultures that were unique (highly expressed) to each age group (as shown in colour legend). Blue text highlights those that are highly expressed in mock compared to SARS-CoV-2 infection conditions and black text enriched with infection. **(f)** Total numbers of annotated airway epithelial cells in all mock infected vs all SARS-CoV-2 infected datasets in respect to age (data shown in ratio of cell numbers/per 1000 cells in age dataset). **(g)** Dot plots showing the log-fold change in SARS-CoV-2 versus mock at different time points for all cell types in different age groups. Calculation based upon absolute cell numbers with the average fold-change indicated by colour. The dot size corresponds to the number of that cell type per 1000 cells from each condition. **(h)** Uniform manifold approximation and projection (UMAP) representation of the results from Milo differential abundance testing. Nodes are neighbourhoods, coloured by their log fold change when comparing SARS-CoV-2 infected versus mock conditions in adult samples. Non-significant DA neighbourhoods at FDR 10% are coloured grey and significant DA neighbourhoods at FDR 10% are coloured with increased log fold change in red and decreased log fold change in blue. Node sizes correspond to the number of cells in a neighbourhood. The layout of nodes is determined by the position of the neighbourhood index cell in the UMAP. **(i)** Palantir inferred pseudotime probabilities of cycling basal cells differentiating into Ciliated 1, Basaloid-like 2 or Goblet 2 inflammatory cells

**Extended Figure 6. Paediatric Goblet 2 inflammatory cells and viral truncation in response to IFN signalling**

**(a)** Response to interferon-gamma in AECs. GO term gene signatures scores for the term: response to interferon-gamma (GO:0034341) across cell types. Scores were calculated with Scanpy as the average expression of the signature genes subtracted with the average expression of randomly selected genes from bins of corresponding expression values. **(b)** Gene Set Enrichment Analysis (GSEA) indicating enriched gene ontology terms for Goblet 2 inflammatory cells obtained using ShinyGo. **(c)** Volcano plot showing differential expressed proteins of the apical secretome between mock and SARS-CoV-2 infected cultures that were unique (highly expressed) in the paediatric cohort. Blue highlights those that are highly expressed in mock compared to SARS-CoV-2 infection conditions and black enriched with infection. **(d)** GSEA for expression of apical secretome genes of paediatric cells at 72h p.i with SARS-CoV-2 obtained using ShinyGo. Coverage plots of viral reads aligned to SARS-CoV-2 genome in each cell type in **(e)** paediatric, **(f)** adult **(g)** and elderly infected NECs grouped across all time points. Coverage plots of viral reads aligned to SARS-CoV-2 genome for all cell types, shown by age group, both **(h)** with and **(i)** without spiked-in primer grouped across all timepoints (see methods for more details of primer) and at **(j)** 72h p.i time points. Viral reads for all coverage plots are shown in 100 nucleotide (nt) bins normalised per 5,000 cells. **(k)** Histogram displaying frequency and position of genomic mutations in SARS-CoV-2 consensus sequences from 72h p.i with SARS-CoV-2 (n= P5, A5, E5). Bin size is 1000 bases (left) and 50 bases (right). Colour blocks indicate the start coordinates of annotated viral genes.

Extended Figure 7. Transmission electron micrographs of Paediatric Goblet 2 inflammatory cells

Transmission electron micrographs of Goblet cells from a paediatric donor 72h p.i. with SARS-CoV-2. The panels show different magnifications, the scale bar is given on the left (top panels) or above (bottom panels) of each image. Viral particles are indicated with white arrows.

**Extended Figure 8. Elderly basaloid-like 2 cells drive ITGB6 production and enhance viral pathogenesis**

**(a)** Abundance of ITGB6 protein in the apical secretome of mock or SARS-CoV-2 infected NECs at 72h p.i. for each age group (n=P5, A5, E5). As detected using mass spectrometry.

**(b)** Exemplar Western blot showing vimentin (vim) in the elderly sample 72h p.i. with mock (-) or SARS-CoV-2 (+). GAPDH is the loading control, E-Cadherin is also given for reference.

**(c)** Orthogonal views of the z-stacks showing the location of ITGB6 (green) and KRT5+ve cells (white) from exemplar air–liquid interface cultures, counterstained for F-actin (phalloidin, red) and cell nucleus (DAPI, blue). **(d)** Representative maximum intensity projections (left) and orthogonal sections (right) of immunofluorescence z-stacks of Basaloid-like 2 cell markers ITGB6 (green), KRT5 (white), spike (magenta) counterstained for F-actin (phalloidin; red) and cell nucleus (DAPI, blue) in 72h p.i NECs (mock top, infected bottom). **(e-g)** Further, example immunofluorescence images of Basaloid-like 2 cell markers in 72h p.i. with SARS-CoV-2. Markers and respective counterstain colour are indicated. **(h)** Transmission electron micrograph of non-basal KRT5+ve epithelial cell (white arrow) location within an NEC culture at 72h p.i with SARS-CoV-2. Scale is on the right of the image.

**Extended Figure 9. Wound repair promotes SARS-CoV-2 viral replication**

**(a)** Dotplot visualisation showing viral entry genes for all cell types, with fraction of expressing cells and average expression within each cell type indicated by dot size and colour, respectively. Appended bar graphs indicate absolute cell numbers per cell type. **(b)** Example immunofluorescence orthogonal sections of Basaloid-like 2 cell markers in unwounded or wounded NECs 24h p.i with mock or SARS-CoV-2 infection. F-actin (Phalloidin; red), KRT5 (white), ITGB6 (cyan) and DAPI (blue). **(c)** Mean KRT5 fluorescence signal (RFU) around wound area in different age groups. From maximal projections of fixed NECs without (-) and with (+) wounds, 24 h post wounding. Subject to mixed-effects analysis with sidak’s multiple comparisons test (n=P2, A2, E2) (n=6). **(d)** Example immunofluorescence orthogonal sections of Basaloid-like 2 cell markers in non-wounded or wounded NECs 24h p.i. with mock or SARS-CoV-2 infection from a paediatric and elderly donor. F-actin (Phalloidin; white), Vimentin (VIM) (yellow) and DAPI (blue). **(e)** Mean Vimentin fluorescence signal (RFU) around wound area in different age groups. From maximal projections of fixed AECs without (-) and with (+) wounds, 24 h post wounding. Subject to mixed-effects analysis with sidak’s multiple comparisons test (n=P2, A2, E2) (n=6). **(f)** Maximum intensity projection image of a NEC culture, 24h post wound and 24h p.i. with SARS-CoV-2. Stained for F-actin (Phalloidin; red), ITGB6 (green), SARS-CoV-2 Spike protein (magenta), KRT5 (white) and composite with DAPI (blue). **(g)** Mean ITGB6 fluorescence signal (RFU) around wound area in different age groups. From maximal projections of fixed NECs without (-) and with (+) wounds, 24 h post wounding. Subject to mixed-effects analysis with sidak’s multiple comparisons test (n=P3, A3-4, E3-2).**(h)** Example of wound healing images taken of NECs from different age groups and with mock or SARS-CoV-2 infection. Acquired by light microscopy over 24 hours. The scale bar (bottom right) represents 1 mm. **(i)** Representative Immunofluorescent images from 72h p.i NECs with SARS-CoV-2 without (top) and with (bottom) wounding stained for ITGB6 (cyan) and dsRNA (yellow). Percentage area covered (right) with ITGB6+ve signal or dsRNA+ve signal from maximum intensity projections of fixed NECs using threshold analysis (red) in ImageJ, the percentage coverage is given at the bottom right of each image.

## References

1. Saatci, D. et al. Association Between Race and COVID-19 Outcomes Among 2.6 Million Children in England. JAMA Pediatr. 175, 928–938 (2021).

2. Lu, X. et al. SARS-CoV-2 Infection in Children. N. Engl. J. Med. 382, 1663–1665 (2020).

3. Chou, J., Thomas, P. G. & Randolph, A. G. Immunology of SARS-CoV-2 infection in children. Nat. Immunol. 23, 177–185 (2022).

4. Götzinger, F. et al. COVID-19 in children and adolescents in Europe: a multinational, multicentre cohort study. The Lancet. Child & adolescent health 4, 653–661 (2020).

5. Du, Y. et al. Clinical Features of 85 Fatal Cases of COVID-19 from Wuhan. A Retrospective Observational Study. Am. J. Respir. Crit. Care Med. 201, 1372–1379 (2020).

6. Yanez, N. D., David Yanez, N., Weiss, N. S., Romand, J.-A. & Treggiari, M. M. COVID- 19 mortality risk for older men and women. BMC Public Health vol. 20 Preprint at https://doi.org/10.1186/s12889-020-09826-8 (2020).

7. Kang, S. J. & Jung, S. I. Age-Related Morbidity and Mortality among Patients with COVID-19. Infect Chemother 52, 154–164 (2020).

8. Sungnak, W. et al. SARS-CoV-2 entry factors are highly expressed in nasal epithelial cells together with innate immune genes. Nat. Med. 26, 681–687 (2020).

9. Bridges, J. P., Vladar, E. K., Huang, H. & Mason, R. J. Respiratory epithelial cell responses to SARS-CoV-2 in COVID-19. Thorax vol. 77 203–209 Preprint at https://doi.org/10.1136/thoraxjnl-2021-217561 (2022).

10. Aguiar, J. A. et al. Gene expression and protein profiling of candidate SARS-CoV-2 receptors in human airway epithelial cells and lung tissue. Eur. Respir. J. 56, (2020).

11. Ahn, J. H. et al. Nasal ciliated cells are primary targets for SARS-CoV-2 replication in the early stage of COVID-19. J. Clin. Invest. 131, (2021).

12. Zhou, M. et al. Proteomics profiling of epithelium-derived exosomes from nasal polyps revealed signaling functions affecting cellular proliferation. Respiratory Medicine vol. 162 105871 Preprint at https://doi.org/10.1016/j.rmed.2020.105871 (2020).

13. Chen, J. et al. Nonmuscle myosin heavy chain IIA facilitates SARS-CoV-2 infection in human pulmonary cells. Proc. Natl. Acad. Sci. U. S. A. 118, (2021).

14. Huang, J. et al. SARS-CoV-2 Infection of Pluripotent Stem Cell-Derived Human Lung Alveolar Type 2 Cells Elicits a Rapid Epithelial-Intrinsic Inflammatory Response. Cell Stem Cell 27, 962–973.e7 (2020).

15. Hou, Y. et al. Aging-related cell type-specific pathophysiologic immune responses that exacerbate disease severity in aged COVID-19 patients. Aging Cell 21, e13544 (2022).

16. Chu, H. et al. Comparative tropism, replication kinetics, and cell damage profiling of SARS-CoV-2 and SARS-CoV with implications for clinical manifestations, transmissibility, and laboratory studies of COVID-19: an observational study. Lancet Microbe 1, e14–e23 (2020).

17. Bunyavanich, S., Do, A. & Vicencio, A. Nasal Gene Expression of Angiotensin-Converting Enzyme 2 in Children and Adults. JAMA vol. 323 2427 Preprint at https://doi.org/10.1001/jama.2020.8707 (2020).

18. Muus, C. et al. Single-cell meta-analysis of SARS-CoV-2 entry genes across tissues and demographics. Nat. Med. 27, 546–559 (2021).

19. 19. Yoshida, M., et al. Local and systemic responses to SARS-CoV-2 infection in children and adults. Nature vol. 602 321–327 Preprint at https://doi.org/10.1038/s41586-021-04345-x (2022).

20. Steinman, J. B., Lum, F. M., Ho, P. P.-K., Kaminski, N. & Steinman, L. Reduced development of COVID-19 in children reveals molecular checkpoints gating pathogenesis illuminating potential therapeutics. Proc. Natl. Acad. Sci. U. S. A. 117, 24620–24626 (2020).

21. Loske, J. et al. Pre-activated antiviral innate immunity in the upper airways controls early SARS-CoV-2 infection in children. Nat. Biotechnol. 40, 319–324 (2022).

22. Vono, M. et al. Robust innate responses to SARS-CoV-2 in children resolve faster than in adults without compromising adaptive immunity. Cell Rep. 37, 109773 (2021).

23. Filippatos, F., Tatsi, E.-B. & Michos, A. Immune response to SARS-CoV-2 in children: A review of the current knowledge. Pediatr Investig 5, 217–228 (2021).

24. Habermann, A. C. et al. Single-cell RNA sequencing reveals profibrotic roles of distinct epithelial and mesenchymal lineages in pulmonary fibrosis. Sci Adv 6, eaba1972 (2020).

25. Ziegler, C. G. K. et al. Impaired local intrinsic immunity to SARS-CoV-2 infection in severe COVID-19. Cell 184, 4713–4733.e22 (2021).

26. Hoffmann, M. et al. SARS-CoV-2 Cell Entry Depends on ACE2 and TMPRSS2 and Is Blocked by a Clinically Proven Protease Inhibitor. Cell 181, 271–280.e8 (2020).

27. Dann, E., Henderson, N. C., Teichmann, S. A., Morgan, M. D. & Marioni, J. C. Differential abundance testing on single-cell data using k-nearest neighbor graphs. Nat. Biotechnol. 40, 245–253 (2022).

28. Breuss, J. M., Gillett, N., Lu, L., Sheppard, D. & Pytela, R. Restricted distribution of integrin beta 6 mRNA in primate epithelial tissues. Journal of Histochemistry & Cytochemistry vol. 41 1521–1527 Preprint at https://doi.org/10.1177/41.10.8245410 (1993).

29. Haapasalmi, K. et al. Keratinocytes in human wounds express alpha v beta 6 integrin. J. Invest. Dermatol. 106, 42–48 (1996).

30. Munger, J. S. et al. The integrin alpha v beta 6 binds and activates latent TGF beta 1: a mechanism for regulating pulmonary inflammation and fibrosis. Cell 96, 319–328 (1999).

31. Hatton, C. F. et al. Delayed induction of type I and III interferons mediates nasal epithelial cell permissiveness to SARS-CoV-2. Nat. Commun. 12, 7092 (2021).

32. Tapia, K. et al. Defective viral genomes arising in vivo provide critical danger signals for the triggering of lung antiviral immunity. PLoS Pathog. 9, e1003703 (2013).

33. Genoyer, E. & López, C. B. The Impact of Defective Viruses on Infection and Immunity. Annu Rev Virol 6, 547–566 (2019).

34. Zhang, Y. et al. The diverse roles and dynamic rearrangement of vimentin during viral infection. J. Cell Sci. 134, (2020).

35. Zhao, S.-H. et al. Basigin-2 is the predominant basigin isoform that promotes tumor cell migration and invasion and correlates with poor prognosis in epithelial ovarian cancer. J. Transl. Med. 11, 92 (2013).

36. Stewart, C. A. et al. Lung cancer models reveal SARS-CoV-2-induced EMT contributes to COVID-19 pathophysiology. bioRxiv (2021) doi:10.1101/2020.05.28.122291.

37. Aydillo, T. et al. Shedding of Viable SARS-CoV-2 after Immunosuppressive Therapy for Cancer. New England Journal of Medicine vol. 383 2586–2588 Preprint at https://doi.org/10.1056/nejmc2031670 (2020).

38. Koch, C. M. et al. Age-related Differences in the Nasal Mucosal Immune Response to SARS-CoV-2. Am. J. Respir. Cell Mol. Biol. 66, 206–222 (2022).

39. Pierce, C. A. et al. Natural mucosal barriers and COVID-19 in children. JCI Insight 6, (2021).

40. Beucher, G. et al. Bronchial epithelia from adults and children: SARS-CoV-2 spread via syncytia formation and type III interferon infectivity restriction. Proc. Natl. Acad. Sci. U. S. A. 119, e2202370119 (2022).

41. Walker, F. C., Sridhar, P. R. & Baldridge, M. T. Differential roles of interferons in innate responses to mucosal viral infections. Trends Immunol. 42, 1009–1023 (2021).

42. Park, A. & Iwasaki, A. Type I and Type III Interferons - Induction, Signaling, Evasion, and Application to Combat COVID-19. Cell Host Microbe 27, 870–878 (2020).

43. Wang, K. et al. CD147-spike protein is a novel route for SARS-CoV-2 infection to host cells. Signal Transduct Target Ther 5, 283 (2020).

44. Heald-Sargent, T. et al. Age-Related Differences in Nasopharyngeal Severe Acute Respiratory Syndrome Coronavirus 2 (SARS-CoV-2) Levels in Patients With Mild to Moderate Coronavirus Disease 2019 (COVID-19). JAMA Pediatr. 174, 902–903 (2020).

45. Jones, T. C. et al. Estimating infectiousness throughout SARS-CoV-2 infection course. Science 373, (2021).

46. Despres, H. W. et al. Measuring infectious SARS-CoV-2 in clinical samples reveals a higher viral titer:RNA ratio for Delta and Epsilon vs. Alpha variants. Proc. Natl. Acad. Sci. U. S. A. 119, e2116518119 (2022).

47. Mautner, L. et al. Replication kinetics and infectivity of SARS-CoV-2 variants of concern in common cell culture models. Virol. J. 19, 76 (2022).

48. Kim, Y.-I. et al. Age-dependent pathogenic characteristics of SARS-CoV-2 infection in ferrets. Nat. Commun. 13, 21 (2022).

49. Meecham, A. & Marshall, J. F. The ITGB6 gene: its role in experimental and clinical biology. Gene **763S**, 100023 (2020).

50. Annes, J. P., Rifkin, D. B. & Munger, J. S. The integrin alphaVbeta6 binds and activates latent TGFbeta3. FEBS Lett. 511, 65–68 (2002).

51. Sheppard, D. Functions of pulmonary epithelial integrins: from development to disease. Physiol. Rev. 83, 673–686 (2003).

52. Miyake, S. et al. Beta 1 integrin-mediated interaction with extracellular matrix proteins regulates cytokine gene expression in synovial fluid cells of rheumatoid arthritis patients. J. Exp. Med. 177, 863–868 (1993).

53. de Fougerolles, A. R. et al. Global expression analysis of extracellular matrix-integrin interactions in monocytes. Immunity 13, 749–758 (2000).

54. Marconi, G. D. et al. Epithelial-Mesenchymal Transition (EMT): The Type-2 EMT in Wound Healing, Tissue Regeneration and Organ Fibrosis. Cells 10, (2021).

55. Vareille, M., Kieninger, E., Edwards, M. R. & Regamey, N. The airway epithelium: soldier in the fight against respiratory viruses. Clin. Microbiol. Rev. 24, 210–229 (2011).

56. Ruan, T. et al. H1N1 Influenza Virus Cross-Activates Gli1 to Disrupt the Intercellular Junctions of Alveolar Epithelial Cells. Cell Rep. 31, 107801 (2020).

57. Stewart, C. A. et al. Lung Cancer Models Reveal Severe Acute Respiratory Syndrome Coronavirus 2–Induced Epithelial-to-Mesenchymal Transition Contributes to Coronavirus Disease 2019 Pathophysiology. Journal of Thoracic Oncology vol. 16 1821–1839 Preprint at https://doi.org/10.1016/j.jtho.2021.07.002 (2021).

58. Pandolfi, L. et al. Neutrophil extracellular traps induce the epithelial-mesenchymal transition: implications in post-COVID-19 fibrosis. Preprint at https://doi.org/10.1101/2020.11.09.374769.

59. Downes, D. J. et al. Identification of LZTFL1 as a candidate effector gene at a COVID- 19 risk locus. Nature Genetics vol. 53 1606–1615 Preprint at https://doi.org/10.1038/s41588-021-00955-3 (2021).

60. Kalluri, R. & Weinberg, R. A. The basics of epithelial-mesenchymal transition. J. Clin. Invest. 119, 1420–1428 (2009).

61. Morrison, C. B. et al. SARS-CoV-2 infection of airway cells causes intense viral and cell shedding, two spreading mechanisms affected by IL-13. Proc. Natl. Acad. Sci. U. S. A. 119, e2119680119 (2022).

62. Taddei, M. L., Giannoni, E., Fiaschi, T. & Chiarugi, P. Anoikis: an emerging hallmark in health and diseases. J. Pathol. 226, 380–393 (2012).

63. Matarrese, P. et al. The HIV-1 vpr protein induces anoikis-resistance by modulating cell adhesion process and microfilament system assembly. Cell Death Differ. 7, 25–36 (2000).

64. Kakavandi, E., Shahbahrami, R., Goudarzi, H., Eslami, G. & Faghihloo, E. Anoikis resistance and oncoviruses. J. Cell. Biochem. 119, 2484–2491 (2018).

65. Sadri Nahand, J., et al. Cell death pathways and viruses: Role of microRNAs. Mol. Ther. Nucleic Acids 24, 487–511 (2021).

66. Cao, Z., Livas, T. & Kyprianou, N. Anoikis and EMT: Lethal ‘Liaisons’ during Cancer Progression. Crit. Rev. Oncog. 21, 155–168 (2016).

67. Lu, Y., Liu, D. X. & Tam, J. P. Lipid rafts are involved in SARS-CoV entry into Vero E6 cells. Biochem. Biophys. Res. Commun. 369, 344–349 (2008).

68. Salanueva, I. J., Cerezo, A., Guadamillas, M. C. & del Pozo, M. A. Integrin regulation of caveolin function. J. Cell. Mol. Med. 11, 969–980 (2007).

69. Bergelson, B. A. et al. Prediction of risk for hemodynamic compromise during percutaneous transluminal coronary angioplasty. Am. J. Cardiol. 70, 1540–1545 (1992).

70. Wickham, T. J., Filardo, E. J., Cheresh, D. A. & Nemerow, G. R. Integrin alpha v beta 5 selectively promotes adenovirus mediated cell membrane permeabilization. J. Cell Biol. 127, 257–264 (1994).

71. 71. Embong, A. K., Duffney, P., Thatcher, T. H. & Sime, P. J. Cigarette Smoke Increases Uptake of Influenza A Virus by Lung Epithelial Cells by Increasing Expression of Caveolin-1. B101. ADVANCES IN COPD PATHOGENESIS Preprint at https://doi.org/10.1164/ajrccm-conference.2019.199.1_meetingabstracts.a4051 (2019).

72. Simons, P. et al. Integrin activation is an essential component of SARS-CoV-2 infection. Sci. Rep. 11, 20398 (2021).

73. Butler, C. R. et al. Rapid Expansion of Human Epithelial Stem Cells Suitable for Airway Tissue Engineering. Am. J. Respir. Crit. Care Med. 194, 156–168 (2016).

74. Bliss, E., Heywood, W. E., Benatti, M., Sebire, N. J. & Mills, K. An optimised method for the proteomic profiling of full thickness human skin. Biol. Proced. Online 18, 15 (2016).

75. Schindelin, J., et al. Fiji: An open-source platform for biological-image analysis. Nature Methods vol. 9 676–682 Preprint at https://doi.org/10.1038/nmeth.2019 (2012).

76. B Worlock, K., Yoshida, M., Meyer, K. & Z. Nikolić bronchial and tracheal brushings withcold-active protease for single-cell RNA-seq v1. (2021) doi:10.17504/protocols.io.btpunmnw.

77. Young, M. D. & Behjati, S. SoupX removes ambient RNA contamination from droplet-based single-cell RNA sequencing data. Gigascience 9, (2020).

78. Lun, A. T. L. et al. EmptyDrops: distinguishing cells from empty droplets in droplet-based single-cell RNA sequencing data. Genome Biol. 20, 63 (2019).

79. Wolock, S. L., Lopez, R. & Klein, A. M. Scrublet: Computational Identification of Cell Doublets in Single-Cell Transcriptomic Data. Cell Syst 8, 281–291.e9 (2019).

80. Popescu, D.-M. et al. Decoding human fetal liver haematopoiesis. Nature 574, 365–371 (2019).

81. Pijuan-Sala, B. et al. A single-cell molecular map of mouse gastrulation and early organogenesis. Nature 566, 490–495 (2019).

82. Heaton, H. et al. Souporcell: robust clustering of single-cell RNA-seq data by genotype without reference genotypes. Nat. Methods 17, 615–620 (2020).

83. Polański, K. et al. BBKNN: fast batch alignment of single cell transcriptomes. Bioinformatics 36, 964–965 (2020).

84. Traag, V. A., Waltman, L. & van Eck, N. J. From Louvain to Leiden: guaranteeing well-connected communities. Sci. Rep. 9, 5233 (2019).

85. Wolf, F. A., Angerer, P. & Theis, F. J. SCANPY: large-scale single-cell gene expression data analysis. Genome Biol. 19, 15 (2018).

86. Domínguez Conde, C., et al. Cross-tissue immune cell analysis reveals tissue-specific features in humans. Science 376, eabl5197 (2022).

87. Trapnell, C. et al. The dynamics and regulators of cell fate decisions are revealed by pseudotemporal ordering of single cells. Nat. Biotechnol. 32, 381–386 (2014).

88. Qiu, X. et al. Single-cell mRNA quantification and differential analysis with Census. Nat. Methods 14, 309–315 (2017).

89. Setty, M. et al. Characterization of cell fate probabilities in single-cell data with Palantir. Nat. Biotechnol. 37, 451–460 (2019).

90. Ge, S. X., Jung, D. & Yao, R. ShinyGO: a graphical gene-set enrichment tool for animals and plants. Bioinformatics 36, 2628–2629 (2020).

