## Extended Figure for "The emergence of goblet inflammatory or ITGB6^hi^ nasal progenitor cells determines age-associated SARS-CoV-2 pathogenesis" (from another version of this preprint: version 1, 2023-01-17)

### Extended Figure 1

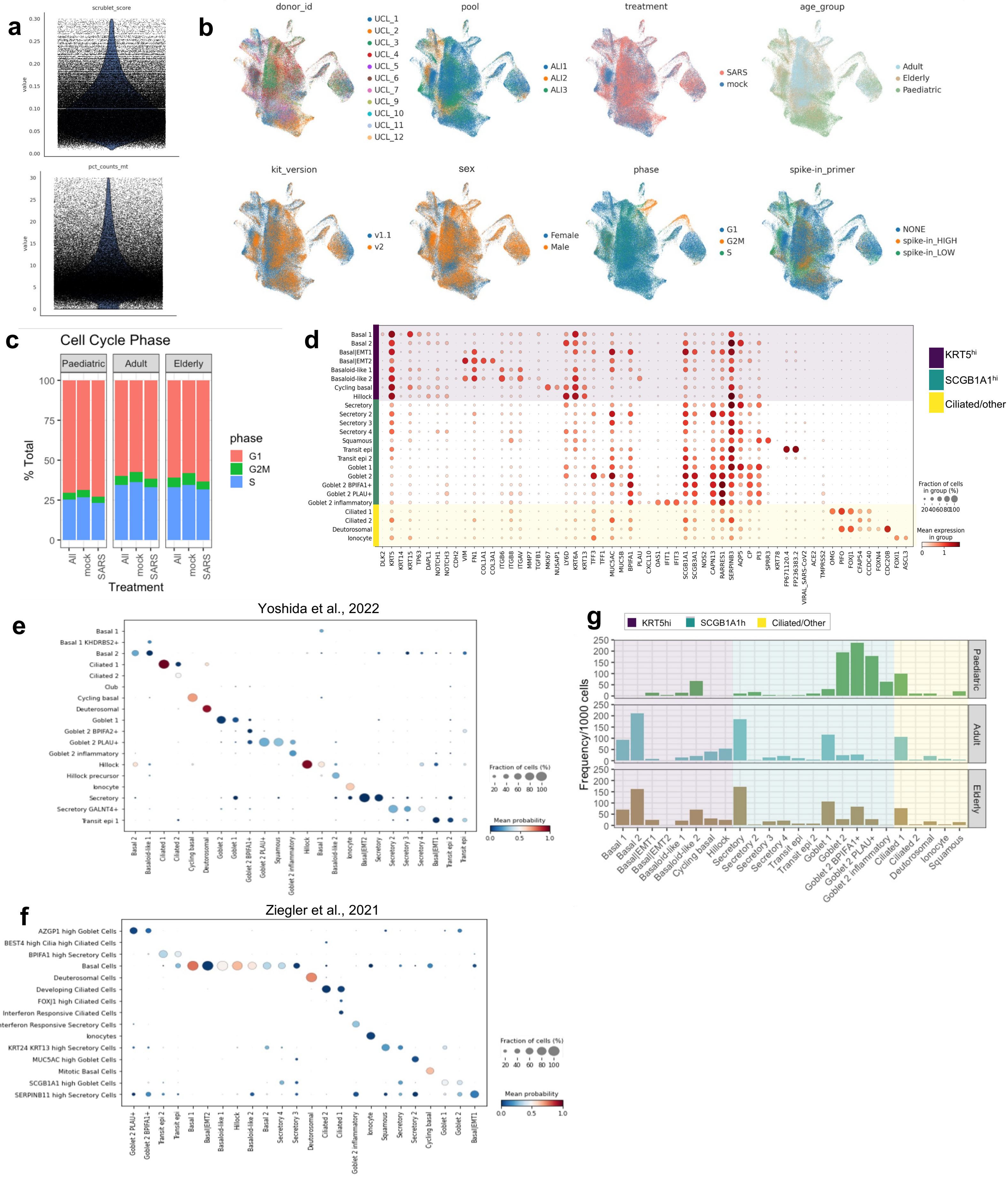

### Extended Figure 2

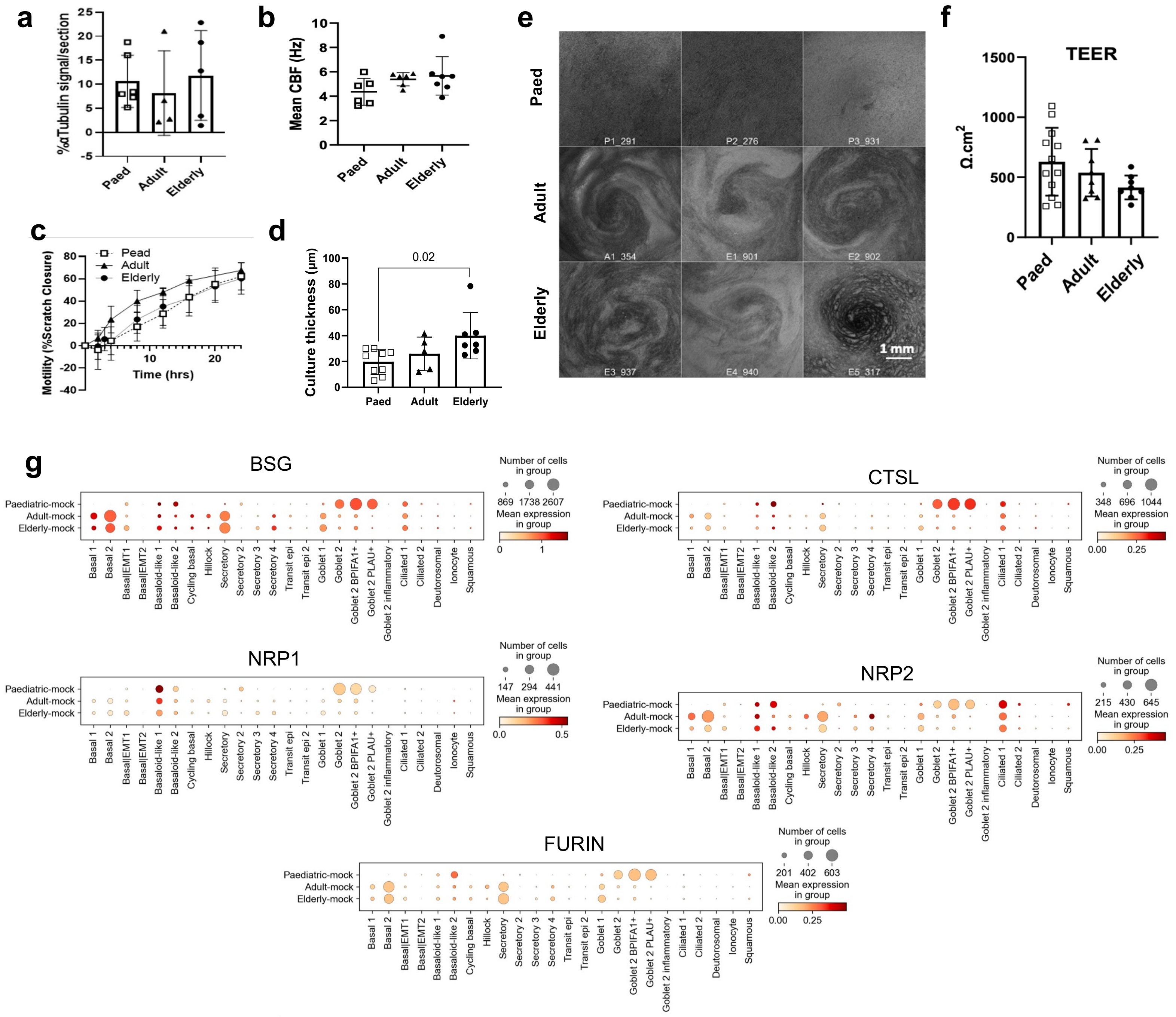

### Extended Figure 3

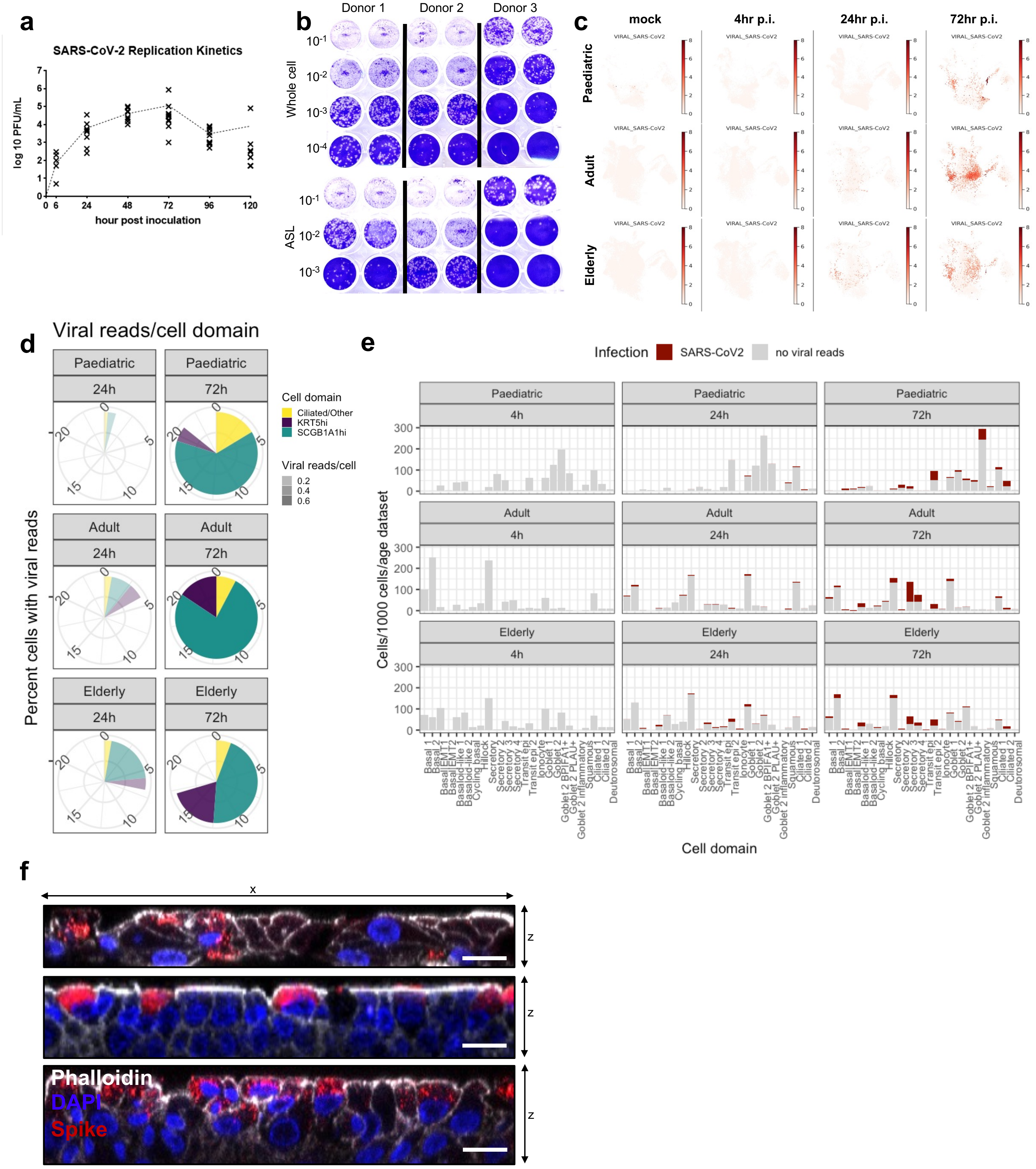

### Extended Figure 4

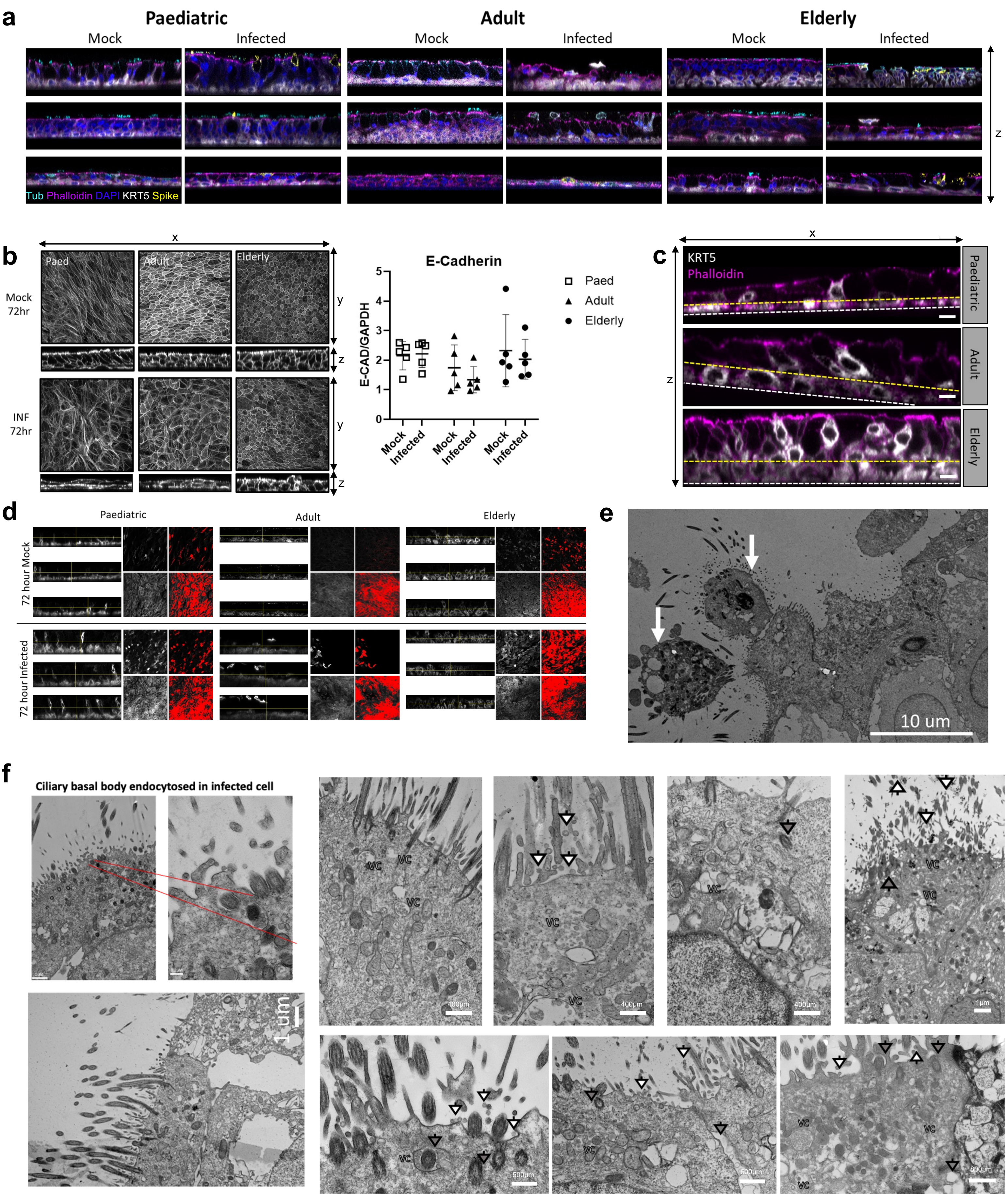

#### Extended Figure 5

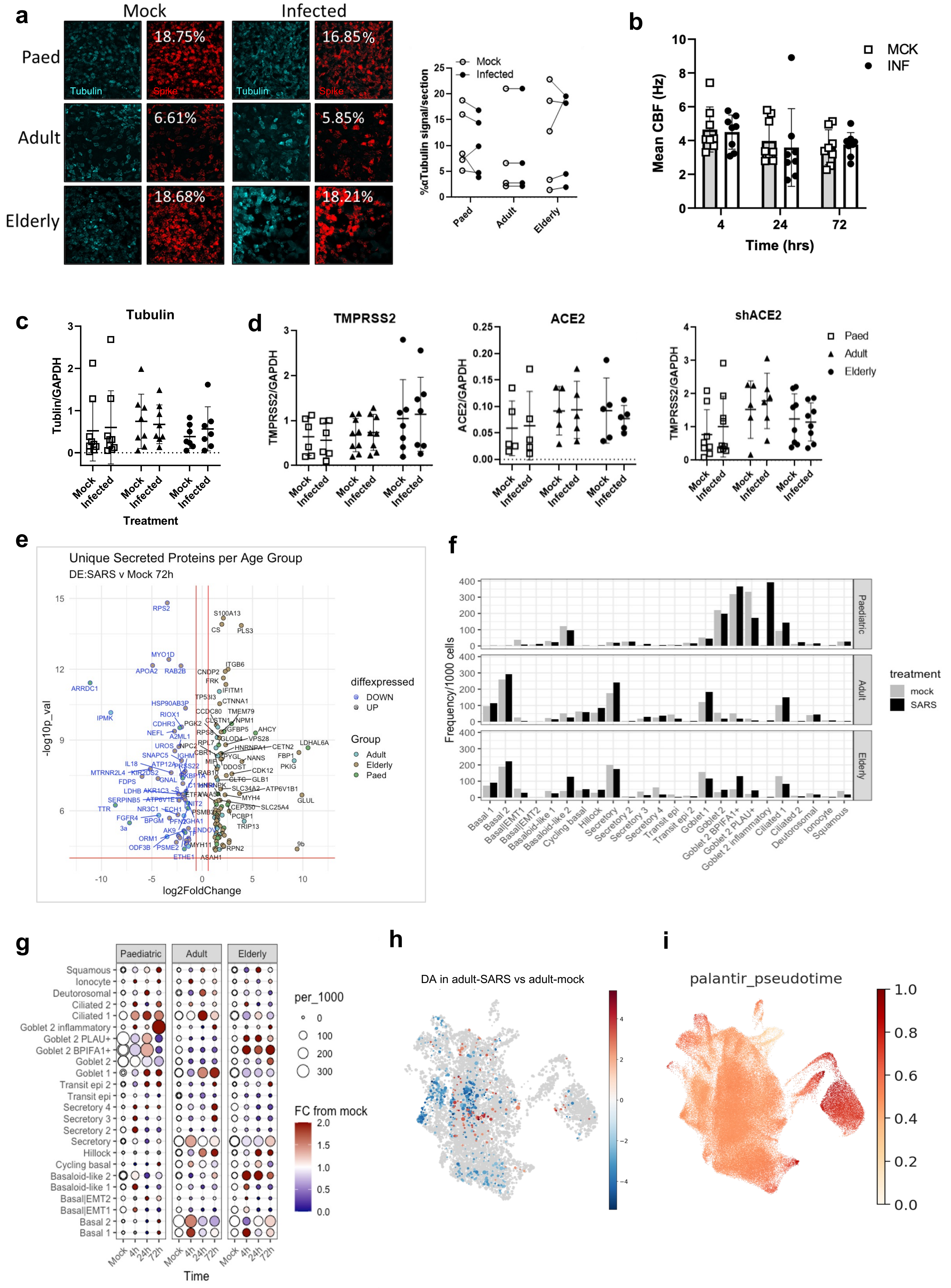

#### Extended Figure 6

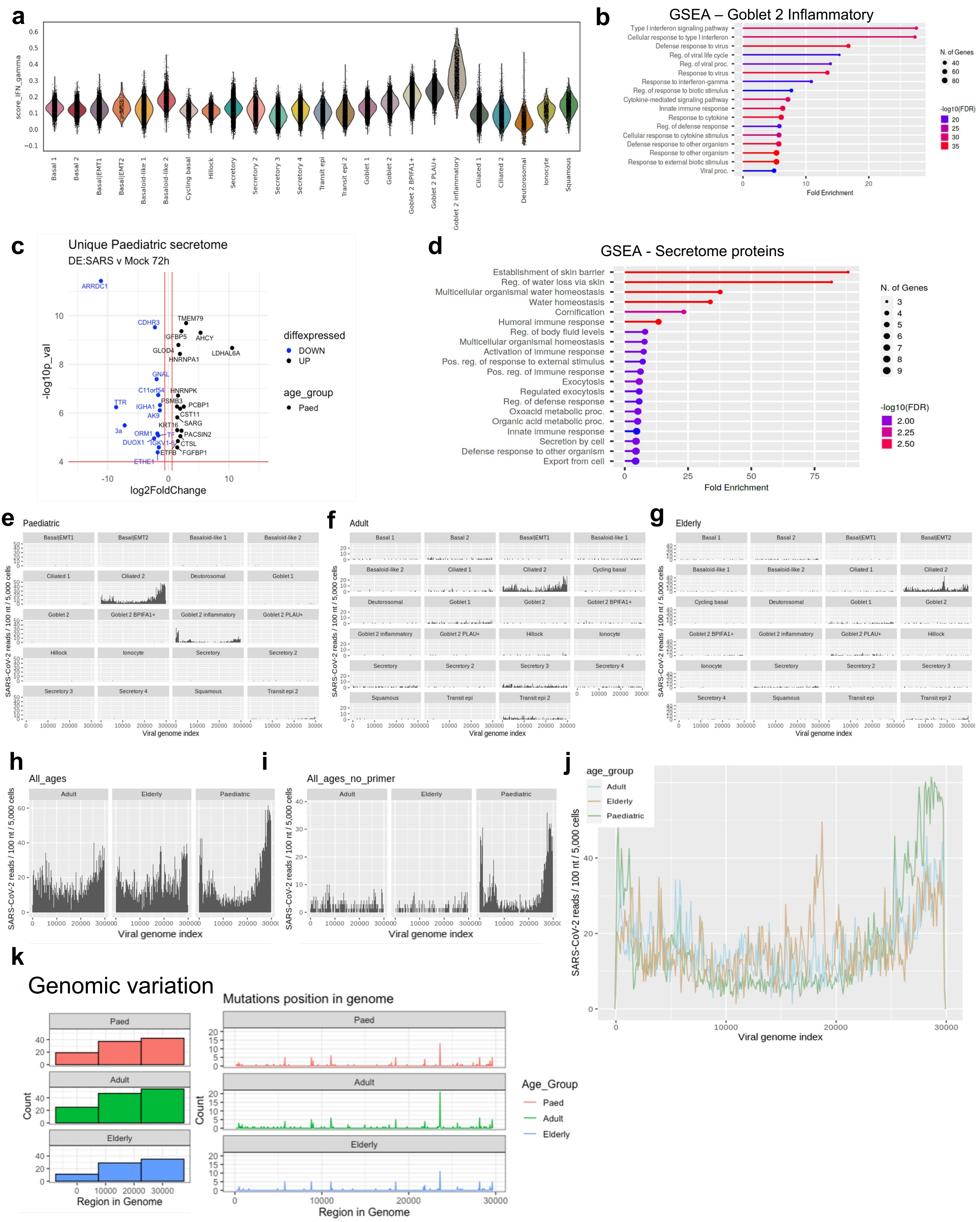

### Extended Figure 7

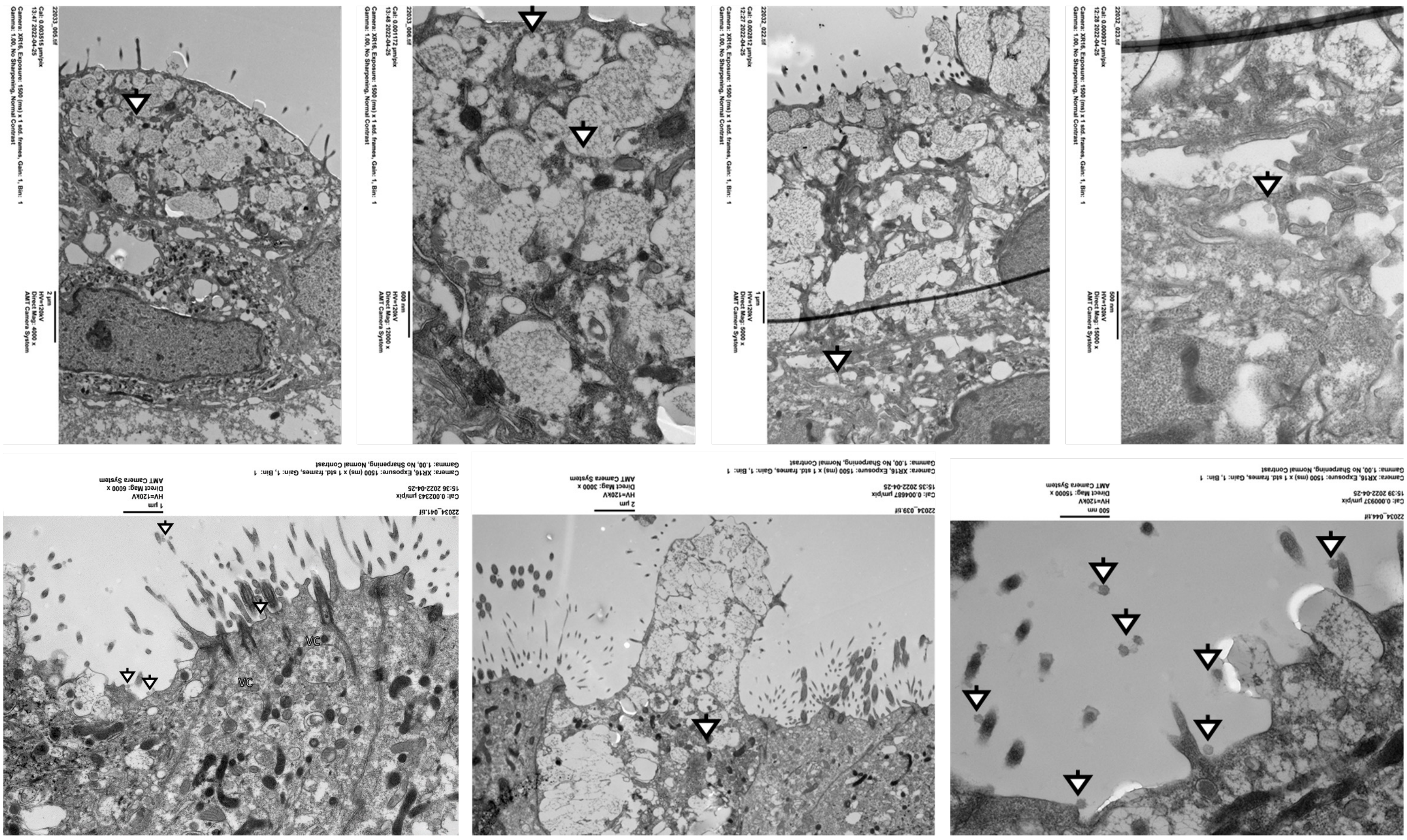

### Extended Figure 8

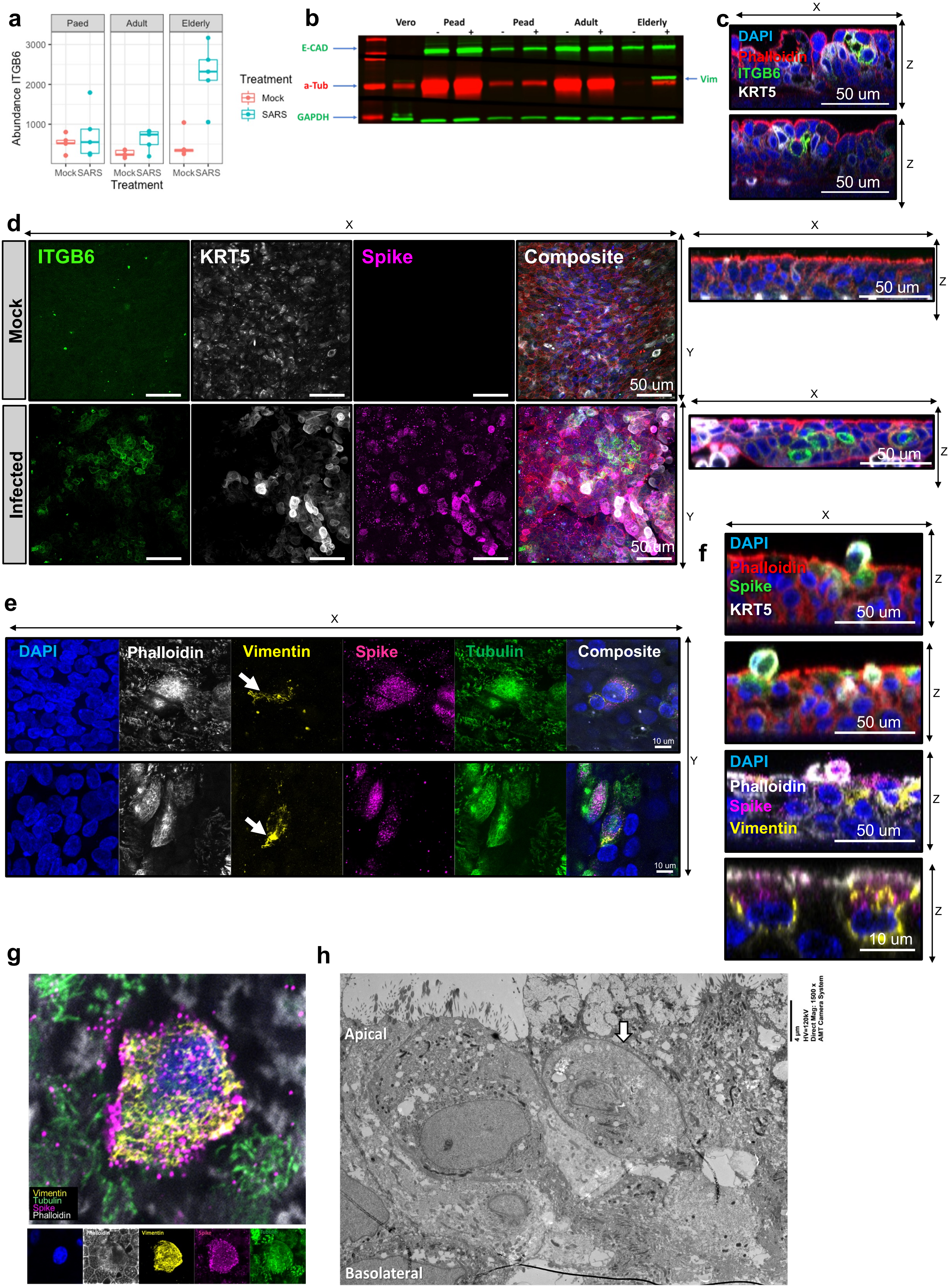

### Extended Figure 9

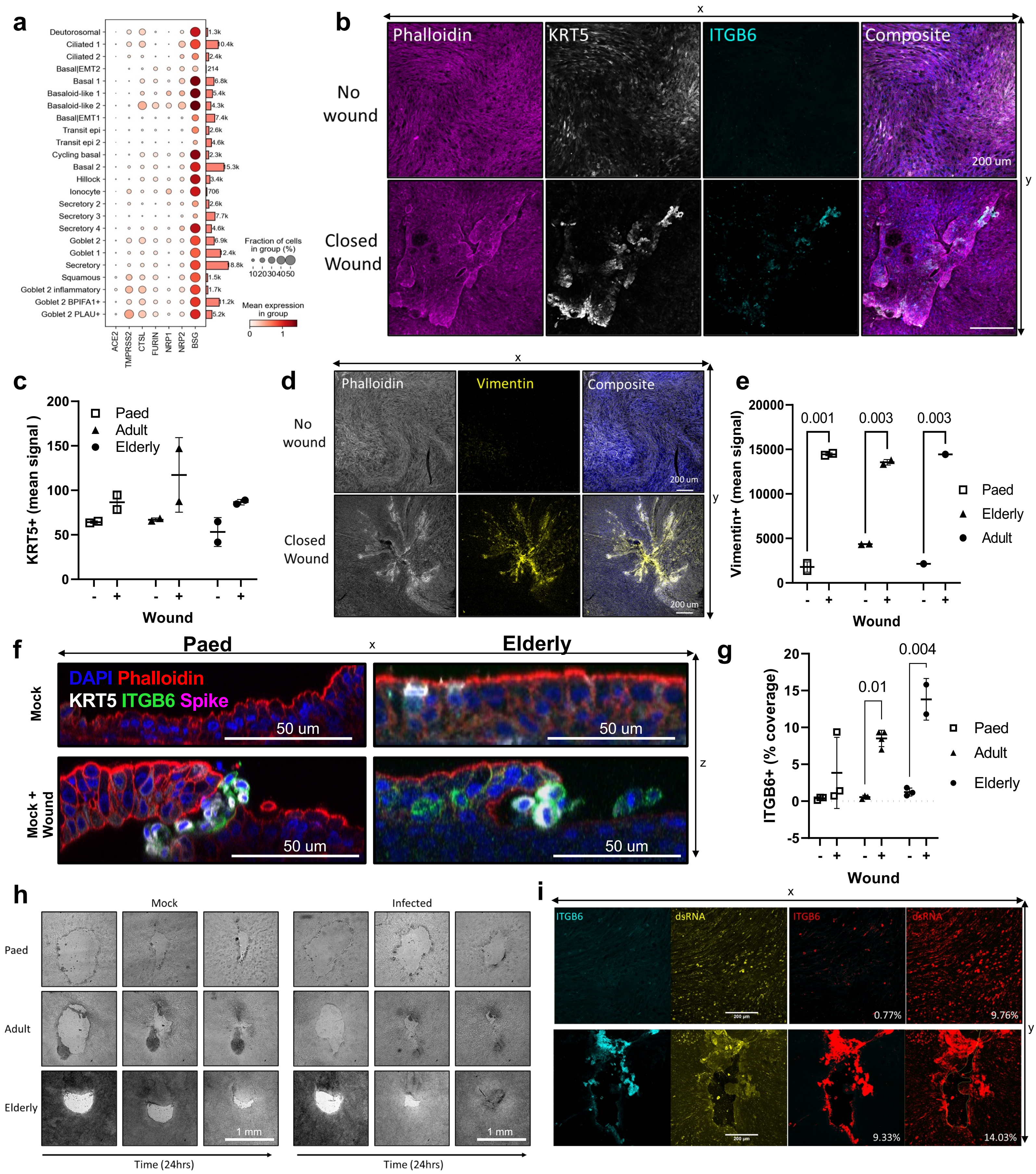
